# Commonness and rarity of pollinators: species abundance and diversity in a hyperdiverse Mediterranean assemblage

**DOI:** 10.64898/2026.02.03.703540

**Authors:** Carlos M. Herrera

## Abstract

This paper presents a comprehensive analysis of patterns of species abundance and diversity in a hyperdiverse insect pollinator assemblage from Mediterranean montane habitats of southeastern Spain. Data on pollinator visitation to flowers of the community of entomophilous plants (288 species) were gathered over a 29-year period, and ∼95% of individual pollinators recorded were identified to species, totalling 46,401 individuals in 845 species. Shapes of species abundance distributions (SADs) were virtually identical at the regional (*N* = 56 sites) and local (one intensively studied site) scales, remained invariant over the study period, and were best predicted by the log-series distribution. Pollinator diversity estimates corresponding to the first three Hill numbers (Species richness, Shannon diversity and Simpson diversity; ^0^D, ^1^D and ^2^D, respectively) were obtained for each plant species x site x year combinations (*N* = 472 "sampling occasions"). Pollinator diversity measures varied widely among plant species; their frequency distributions were unimodal and strongly right-skewed; and interspecific variation was related to plant phylogeny, floral features and pollinator visitation rates. Pollinator diversity of individual plant species depended on habitat type, with dolomitic outcrops, rock cliffs and forest interior harboring the least diverse pollinator assemblages. Variation of ^0^D, ^1^D and ^2^D among habitats and years was largely decoupled, revealing a complex patterning of pollinator species richness and dominance. Estimated proportions of undetected pollinator diversity ("dark diversity") varied with insect order (lowest for Lepidoptera, highest for Diptera). The adoption of community ecology tools for assessing pollinator diversity improves our ability to elucidate pollinator responses to natural and anthropogenic environmental change, and permits hitherto unexplored questions in pollination ecology.

## INTRODUCTION

The notion of "pollinator diversity" is a transversal thread that pieces together all facets of research on animal pollination (Ollerton 2017; Loy & Brosi 2022), as it is variously related to ecosystem service, ecological functionality, pollination success, plant specialization, crop production, anthropogenic disturbance, global change and biodiversity conservation (Kevan et al. 1997; Herrera 2005; Gómez et al. 2007; Fründ et al. 2010; Davila et al. 2012; Pérez-Méndez et al. 2020; Loy & Brosi 2022; Pérez-Gómez et al. 2024). In the majority of instances, however, when it is applied to animal pollinators in pollination specific research the term "diversity" is closer to its vernacular meaning (a synonym for variety or assortment of kinds) than to the species diversity concept rooted in community and statistical ecology. The latter has historically combined several theoretical concepts including species richness, evenness, dominance, and species abundance distributions (Margalef 1957; Williams 1964; Whittaker 1965; Hill 1973; May 1975; Magurran 2004; Jost 2006, 2010). Pollination studies have only infrequently implemented these concepts (but see, e.g., Tepedino & Stanton 1981; Kato et al. 1990; Kevan et al. 1997; Galloni et al. 2008; Norfolk et al. 2015; Pérez-Gómez et al. 2024; Rezende et al. 2025), some implications of which are highlighted in the following section.

### "Ecologically-informed" *vs.* "Naïve" pollinator diversity

As stressed by Hill (1973, p. 427), "the purpose of determining diversity by a numerical index is […] to provide a means of comparison". Half a century later comparisons of pollinator diversity are frequently based on numbers of pollinator species taken at face value ("naïve" diversity), without attention to sample completeness, sampling biases, or the fact that statistical properties of species richness are much worse than those of any other common diversity measure (Chao & Jost 2012; see, e.g., Appendix 2 to Ollerton 2017 for pertinent examples and literature references). Valid comparisons of pollinator diversity require some quantitative estimation of the impact of incomplete sampling of multispecies assemblages, which in turn requires elucidating the statistical properties of species abundance distributions and the estimation of different measures of species diversity which place variable emphasis on rare and abundant species (Hill 1973; Magurran 2004; Gotelli & Chao 2013). Since species abundance distributions are tightly intertwined with the concepts inherent to "ecologically-informed" diversity (Whittaker 1965; May 1975; Magurran 2004), robust comparisons of pollinator diversity across populations, species, habitats or years should consider how the frequencies of different pollinator species are distributed. In practice, however, it remains essentially unknown the degree to which species abundance distributions in pollinator assemblages conform to the various statistical distributions which have been proposed for other multispecies assemblages (Williams 1964; Magurran 2004; Ulrich et al. 2010; Baldridge et al. 2016; Su 2018; for a limited review for pollinator assemblages see Table 2 of Herrera 1989). In addition, information on pollinator species abundances and sampling properties of pollinator assemblages are essential to estimate the frequency and taxonomic composition of the rarest, usually undetected pollinators ("dark diversity" or "phantom species"; Pärtel et al. 2011; Beck et al. 2018), whose functional importance and perhaps higher extinction risk are being increasingly recognized (Liu et al. 2022; Simpson et al. 2022; Genung et al. 2023).

Statistical properties of pollinator species abundance distributions are expected to influence plant-pollinator systems, as suggested by synthetic papers combining pollination data with other ecosystem functions or using pollination data to make broader conceptual points (Roswell et al. 2021, 2023). Naïve pollinator diversity measures that disregard information on the dominance/evenness of species will miss important aspects of the eco-evolutionary dynamics linking plants and pollinators (Galloni et al. 2008; Winfree et al. 2014, 2015). For example, the more even the relative abundances of species in a pollinator assemblage, the less likely that some particularly abundant pollinator(s) exerts a sufficiently differentiated selection on plant traits as to drive pollinator specialization (Herrera 1989). Species abundance distributions can also provide insights on the variable impact of the extinction of pollinator species occupying different positions along the commonness-rarity gradient (Winfree et al. 2014, 2015). Information on species abundance distributions of pollinators is also needed to formulate null models of the stability and evolution of multispecies mutualistic systems and test whether alleged adaptive features of so-called "plant-pollinator networks" (e.g., connectance, nestedness; Vázquez et al. 2009) might actually represent assembly artefacts, or statistical "spandrels" (Maynard et al. 2018), depending more on hitherto little-investigated statistical and sampling properties of species abundance distributions of pollinators than on adaptive eco-evolutionary processes (Blüthgen et al. 2008; Vázquez et al. 2009; Blüthgen 2010; Maynard et al. 2018; Chacoff et al. 2018; Hu et al. 2019; Thomson 2021).

### General objective, approach and questions

Insufficient knowledge of species abundance distributions and ecologically-informed diversity of pollinator assemblages noted above mainly reflect practical limitations arising from pollination studies mostly dealing with small fractions of local or regional plant assemblages, and involving incompletely identified pollinators. For instance, only 55% of pollinator taxa on average were identified to species in the 24 plant-pollinator network examples included in the bipartite R package (Dormann et al. 2008). Generalizations from small-scale studies to local or regional communities of animal-pollinated plants are also seriously limited by biases in taxonomy, phylogenetic position or floral features of the particular plant species chosen for study (Ollerton et al. 2015; Herrera 2020; Adamo et al. 2021), or by biases or distortions introduced by poor taxonomic resolution (Renaud et al. 2020) and insufficiently validated pollinator identifications (Packer et al. 2018).

The central objective of this study is to elucidate patterns of species abundance and ecologically-informed diversity for the whole insect pollinator assemblage of an entire regional community of entomophilous plants. The information on species abundance distributions thus obtained, in combination with application of the community ecology toolbox to an exhaustive plant-pollinator dataset, will also allow to formulate questions that have remained largely unexplored so far in pollination research (enumerated in the next paragraph). To avoid biases resulting from the selection of a small number of plant species or the variable identification difficulties of different pollinator groups (Herrera et al. 2025), pollinators were thoroughly identified to species and the set of plant species encompassed the entire taxonomic and phylogenetic spectrum of animal-pollinated plants in the study region. This study also provides a case example showing the unique value of the "ecology of place" research paradigm (Price & Billick 2010), based on large samples obtained systematically in the same location over long research spans, for an improved understanding of the ecology and evolution of plant-pollinator systems (Herrera 2020; Janzen & Hallwachs 2021).

As noted above, one advantage of the application of the community ecology toolbox to evaluate pollinator species abundance and diversity is the possibility of formulating some questions that have been rarely addressed so far in pollination studies: (1) How many different species of insect pollinators does a plant community have?; (2) Are discrete levels of plant specialization on pollinators actually recognizable in a plant community (i.e., a multimodal distribution of pollinator diversities)?, or alternatively, Is there a continuous, unimodal gradient in pollinator diversity across plant species?; (3) Do evolutionary history or floral features predict interspecific variation in pollinator species diversity?; (4) Are there some relationships between pollinator diversity and ecological factors such as pollinator abundance, habitat type, flowering phenology or long-term rhythms?; (5) Is species richness a meaningful solo descriptor of pollinator diversity or, alternatively, are variations in dominance/evenness strong enough to blur or distort variations in species richness?; and (6) How large can be the undetected, dark pollinator diversity in a hyperdiverse plant-pollinator system such as the one studied here?, or expressed in statistical terms, What is the magnitude of downward biases in empirical diversity estimates arising from incomplete enumeration of a diverse insect pollinator assemblage? Results of this study will provide answers to these questions.

## MATERIALS AND METHODS

### Study area, habitat types and plant species

Data on species composition of pollinators analyzed in this paper were gathered during 1997-2025 (with a gap in 2000-2002) in the southernmost part of the Sierras de Cazorla-Segura-Las Villas Natural Park (Jaén Province, southeastern Spain). A map of sampling sites and photographs of main habitat types can be found in Herrera (2021) and Herrera (2019), respectively. The Sierra de Cazorla mountain range is an outstanding biodiversity hotspot in the Mediterranean Basin characterized by exceptional plant and animal diversity (Médail & Diadema 2009; Gómez Mercado 2011; Ortiz-Sánchez et al. 2023; Pugnaire et al. 2024; Herrera et al. 2025). Pollinator sampling sites (*N* = 56) were distributed altitudinally (range 770-1920 m a. s. l.) over all major vegetation belts of the region and were a superset of the *N* = 42 locations listed in Appendix S1 to Herrera (2021). From lower to higher elevations, major vegetation types sampled included *Quercus rotundifolia*-dominated evergreen forest and tall scrubland; mixed *Pinus* forest with *Acer* and deciduous *Quercus*; and various types of mature *Pinus nigra* forests and open woodlands differing in age, height, understory and tree density. Nine structural habitat types were recognized for some of the analyses presented here: vertical rock cliffs; local disturbances caused by humans, large mammals or natural abiotic processes; sandy or rocky dolomitic outcrops; dwarf mountain scrub dominated by cushion plants; forest edges and large clearings; forest interior; patches of grasslands and meadows on deep soils in relatively flat terrain; tall, dense Mediterranean sclerophyllous forest and scrub; and banks of permanent streams or flooded/damp areas surrounding springs.

Pollinator species composition was assessed for 292 plant species in 185 genera and 49 families, which are a superset of the 191 species studied by Herrera (2021) and include virtually all species of insect-pollinated plants occurring in the Sierra de Cazorla area. Despite efforts to locate workable populations, ∼15 rare entomophilous species could not be finally included in the study because grew in unaccessible locations or it was impractical to meet the preset requirements for pollinator censuses (see below). Asteraceae (55 species), Lamiaceae (26), Fabaceae (21), Brassicaceae (19), Rosaceae (13) and Cistaceae (11) contributed collectively about half of plant species studied. Each plant species was assigned to one of the nine structural habitat types listed above. Species that occurred in more than one habitat were assigned to the one where pollinator censuses were conducted. Four plant species failed to yield any pollinator observation and were excluded from the data (*Dipcadi serotinum*, *Parentucellia latifolia*, *Sarcocapnos baetica*, *Sherardia arvensis*). The full list of the 288 plant species retained for study can be found in Herrera (2026).

### Sampling scheme

Obtaining pollinator data for such a large species set required spanning field work over many years, since pre-established replication rules (see *Pollinator visitation* below) constrained the number of species sampled per flowering season. About two thirds of plant species (*N* = 189) were sampled for pollinators in only one year, and the rest were sampled on ≥2 years.

Pollinator sampling was conducted at a single site in the vast majority of the species (*N* = 265), while 23 species were sampled on two or more sites. Pollinator sampling effort for each plant species is shown in Herrera (2026). Plant species x site x year combinations ("sampling occasions" hereafter) were chosen randomly (Herrera 2019), subject to constraints set by availability of time and study populations. Many plant populations sampled during the first half of the study had declined substantially or gone extinct over 2018-2025 (∼75 species, mostly from cliffs, dolomitic outcrops, dwarf mountain scrub and damp areas), presumably as a consequence of increasing aridity and temperature (Herrera et al. 2023).

### Pollinator visitation

Pollinator visitation data were obtained by applying the same standardized sampling protocol to all plant species (Herrera 2019). The basic sampling unit was a 3-min watch of a flowering patch whose total number of open flowers was also counted before starting the watch ("pollinator census" hereafter). Pollinators visiting some flower in the focal patch during the watching period were identified (see *Pollinator identification* below), and the total number of flowers probed by each visiting individual was recorded. Areal extent and number of open flowers in watched patches were adjusted for each plant species according to flower size and density, so that I could confidently monitor all pollinator activity in the patch from a distance of 1.5-2 m. Counting the number of elemental florets visited by pollinators was unfeasible in species with tiny flowers densely packed into compact inflorescences (e.g., Apiaceae, Asteraceae). In these cases the number of inflorescences available per patch and visited per census were counted rather than individual flowers, and visitation probabilities (see *Data analysis* below) referred to inflorescences. For simplicity I will refer to visitation to single flowers and inflorescences as "flower visitation". In some analyses the two types of visitation units will be treated as levels of the discrete variable "visitation unit" ("single flowers" *vs.* "flower packets"; see *Data analysis* below).

Census replication rules for each species x site x year combination were as described in Herrera (2019). At least 60 censuses spread over three non-consecutive dates should be conducted on ≥20 widely spaced flowering patches with roughly similar flower numbers. On each date patches were censused in random order and censuses were distributed from 0.5-2.5 h past sunrise (depending on season, censuses started earlier in summer) through one hour past noon. About one third of plant species studied did not have flowers available to pollinators in the afternoon, as their corollas wither, close or fall shortly after noon. Earlier studies in the area also showed a marked afternoon decline of insect pollinator activity (Herrera 1990, 1995). Circumstantial evidence suggesting some crepuscular or nocturnal pollination was found for only four species. Long spells of poor weather, logistic problems or destruction of flowering patches by wild mammals sometimes precluded fulfilling the preceding rules for some species x site x year combinations. The number of sampling dates and pollinator censuses for each plant species are shown in Herrera (2026). A total of 40,666 pollinator censuses were carried out on 854 different dates.

### Pollinator identification

Identification of pollinators recorded in censuses relied on one or more of the following: (1) my familiarity with insect pollinators of the study area; (2) comparison of close-up photographs taken during censuses with a large reference collection of specimens from the study area identified by insect taxonomists; and (3) identification by taxonomists of collected specimens or close-up photographs taken during censuses. Insect taxonomists that contributed identifications for this study are listed in the Acknowledgments. Out of a total of 48,407 pollinator individuals recorded in censuses (all plant species and sampling occasions combined), 95.9% were confidently identified to species (i.e., a full Latin binomial was assigned), and these data form the basis for this study. Records of pollinators not identifiable to species (*N* = 2,006) were purged from the data. The proportion of unidentifiable individuals varied widely among major insect orders, increasing in the direction Lepidoptera (0.3% unidentifiable individuals, mostly Micropterigidae), Hymenoptera (2.1%, mostly *Andrena* and *Lasioglossum* bees), Coleoptera (4.1%, mostly Mordellidae) and Diptera (9.0%, mostly Anthomyiidae, Bombyliidae, Empididae and Muscidae).

Close-up photographs of pollinators visiting flowers were routinely taken during censuses using a DSLR digital camera and 105 mm macro lens. In addition to using them for identification purposes and keeping photographic vouchers, the large photographic collection assembled during this study (∼75,000 pictures) served for ascertaining the pollinator status of the species recorded, particularly for those of small body size. Only species visibly contacting anthers or stigmas, or with pollen grains on the body, are included here as pollinators.

Photographs provided evidence for including in the analyses some insects, such as ants, hemipterans or crickets, whose pollinating role is ordinarily dismissed.

I single-handedly checked and curated all insect identifications, hence results are free from inter-observer heterogeneity in pollinator identification training or skills (Austen et al. 2016; Ratnieks et al. 2016). Some intraobserver variability, however, could have affected the data because of progressive improvement in my insect identification skills. To minimize this source of bias, prior to undertaking the analyses reported here I thoroughly reviewed the whole photographic collection of pollinators obtained during the study. Older species identifications were re-evaluated on the basis of my current knowledge and updated if needed. A list of the 845 pollinator species identified is shown in Herrera (2026).

### Data analysis

All statistical analyses reported in this paper were carried out using the R environment (version 4.5.0; R Core Team 2025). Linear mixed-effects models were fitted using the lmer function in the package lme4 (Bates et al. 2015). Function gamm4 from the gamm4 package (Wood & Scheipl 2025) was used for fitting generalized additive mixed-effects models, and the gam function in the mgcv package for fitting nonparametric regressions (Wood 2017).

Model-adjusted, marginal effects of predictors were estimated from fitted mixed-effects models using the predict_response function in the ggeffects package (Lüdecke 2018).

Two main classes of analyses of pollinator species abundance and diversity will be presented. First, community-wide analyses of pollinator census data from all plant species combined (*N* = 288) will be used to produce a synthetic overview of pollinators’ taxonomic composition, species richness and species abundance distributions. And second, relationships between pollinator diversity and plant intrinsic features, ecological correlates of pollinator diversity, and the magnitude of "dark" pollinator diversity, will be examined at plant species resolution after excluding low-quality pollinator composition data as reckoned by "sample coverage" (see below). Point estimates of pollinator diversity obtained on different sampling occasions (plant x site x year combinations) will be used in these analyses. Details specific to each analysis will be given in the pertinent places of *Results*.

### Species abundance distributions

Two different metrics will be considered initially to assess pollinator abundance, namely total number of insect individuals recorded ("visitors") and total number of flowers visited ("visits") by each pollinator species. As comparisons will reveal a close linear correlation between the two metrics, only abundance data based on number of individuals will be used subsequently.

Empirical species abundance distributions were compared with theoretical predictions from five different statistical models. Four of these were Fisher’s log-series, Poisson lognormal, negative binomial and power law distributions, which were chosen following the criteria of Baldridge et al. (2016): have discrete distributions, allow calculation of likelihoods, represent all "families" of species abundance distributions recognized by McGill et al. (2007), and are widely used in the ecological literature. The fifth statistical model considered was Alonso & McKane’s (2004) neutral metacommunity distribution, which was included specifically for testing whether empirical species abundance distributions of pollinators conform to expectations from the neutral theory of ecological diversity (Hubbell 2001).

Models were fitted to empirical data using maximum likelihood estimation as implemented in the function fitsad of the sads package (Prado et al. 2025). The performance of different models to predict observed pollinator species abundances was compared using an information-theoretical framework and likelihood-based model selection criteria (Burnham & Anderson 2002) as implemented in function AICtab of the bbmle package (Bolker et al. 2023). Burnham & Anderson’s (2002) guidelines will be followed for reporting and interpreting results of model selection. To evaluate whether pooling pollinator data from separate locations had some spurious effect on species abundance distribution, separate analyses were carried out at "regional" and "local" scales. The former included the data from all sampling sites combined, while the latter included only data from the intensively studied, 8.5-ha Nava de las Correhuelas site (NC hereafter; see map in Figure 2 of Herrera 2021 for location and distance relative to the rest of sampling locations). To examine whether the form of species abundance distributions remained consistent over the study period, non-overlapping subsets of data were created by splitting the regional and local datasets into five sequential subperiods with equal number of pollinator individuals. Possible differences among insect taxonomic groups in species abundance distributions were also explored by conducting separate analyses for the four major insect orders.

### Pollinator diversity measures

Measures of "Species richness", "Shannon diversity" and "Simpson diversity", meant here as named in the output of the iNEXT package (Hsieh et al. 2016), will be used throughout this paper as estimators of pollinator diversity (Magurran 2004). They represent the Hill numbers of order *q* = 0, 1, 2, which correspond to total number of species, exponential of Shannon’s entropy, and reciprocal of Simpson’s index, respectively, expressed in the common scale of effective number of species present (Hill 1973; Jost 2006). The three measures provide information on the richness and dominance components of biological diversity by differing in their sensitivity to common and rare species (Hill 1973; Jost 2006; Chao et al. 2014). For example, a steep drop in diversity as *q* increases indicates a high degree of dominance in the community (Jost 2006; Chao & Jost 2015). Although the Hill numbers framework has sometimes attracted theoretically-inspired criticism (Ricotta & Feoli 2024; Alroy 2025), it is a tool of proven heuristic value in investigations of plant-pollinator interactions (Norfolk et al. 2015; Roswell et al. 2021, 2023; Pérez-Gómez et al. 2024).

Empirical pollinator diversity measurements and asymptotic predictions obtained by interpolation-extrapolation (Chao & Jost 2012) were computed with the iNEXT function of the iNEXT package (Hsieh et al. 2016). Empirical and predicted diversity estimates for the three diversity measures were closely correlated across sampling occasions (*r* = 0.832, 0.967 and 0.982 for Species richness, Shannon diversity and Simpson diversity, respectively; *N* = 472 sampling occasions, *P* < 2.2E-16). Unless indicated otherwise, only empirical diversity measurements will be reported and analyzed for simplicity. To test whether discrete degrees of plant specialization on pollinators were recognizable in the plant community as a whole, dip multimodality tests (Hartigan & Hartigan 1985) were applied to the frequency distributions of diversity measures. The iNEXT function was also used to estimate sample completeness, or the Good-Turing’s "sample coverage" parameter (SC), which will be used here to evaluate the quality of pollinator composition data for every sampling occasion (SC = "the proportion of the total number of individuals in a community that belong to the species represented in the sample"; Chao & Jost 2012; Hsieh et al. 2016). In absence of reliable species inventories in the study area for most taxonomic groups of pollinators, the number of species that exist in the pollinator community sampled but were not detected ("dark diversity", *sensu* Pärtel et al. 2011) was estimated with two different statistical methods: the interpolation-extrapolation method of Chao et al. (2014) as implemented by Hsieh et al. (2025) in the R package iNEXT, and the Good-Turing method of Chao et al. (2017) as implemented in the online SuperDuplicates application (https://chao.shinyapps.io/SuperDuplicates/; accessed December 2025). The proportion of total existing species that remained undetected was then estimated in each case as (Predicted - Observed) / Predicted.

### Pollinator diversity and plant inherent features

Phylogenetic relationships among plant species were obtained using the package V.PhyloMaker2 and the WP nomenclature standardization system (Jin & Qian 2022). A test of plant phylogenetic signal in the pollinator diversity data (= the tendency for related species to resemble each other more than they resemble species drawn at random from the phylogenetic tree; Blomberg & Garland 2002) was conducted using Pagel’s λ (Münkemüller et al. 2012) and the package phylosignal (Keck et al. 2016).

Two floral features were examined as potential predictors of pollinator diversity: class of floral perianth and type of pollinator "visitation unit" (single flower *vs.* flower packet; see Appendix S2: Fig. S1 in Herrera 2020). Each plant species was scored for these two discrete variables (Herrera 2026). Two perianth classes were recognized, corresponding to open, more or less bowl-shaped, non-restrictive perianths (“open perianth” hereafter, *N* = 116 species), and closed, tubular, sympetalous or otherwise restrictive perianths (“restrictive perianth” hereafter, *N* = 172 species).

## RESULTS

### Community-wide patterns

This section presents an overview of taxonomic composition and observed species richness of the regional pollinator assemblage; evaluates agreement between three pollinator importance metrics; and examines the statistical properties of species abundance distributions. Separate results will be given for analyses at the local and regional scales. Given the close similarity between spatial scales in all aspects of species abundance distributions reported in this section, subsequent diversity analyses will deal only with pollinator diversity at the regional scale.

### The pollinator assemblage

A total of 845 and 583 species of pollinators belonging to eight insect orders were recorded at the regional (*N* = 288 plant species, 56 sites) and local scales (*N* = 93 plant species, NC site), respectively. Pollinator composition at the insect order level is summarized in Table 1. A close agreement existed between the three importance measures considered (percent contribution of each order to total species, individuals and flower visits), and also between spatial scales. Four insect orders (Hymenoptera, Diptera, Coleoptera, Lepidoptera) accounted altogether for >99% of species, individuals and flower visits at both the regional and local scales. Within this dominant group, Hymenoptera and Diptera were most important irrespective of the importance measure chosen, accounting for more than two thirds of all species, individuals and flowers visited.

**Table 1.** Taxonomic composition of insect pollinators at the regional and local spatial scales. Relative importance of each insect order was assessed by the proportional contribution to total species, individuals and flower visits.

| Insect order | Regional scale<br>(*N* = 845 insect species, 46,401 individuals, 142,177 flower visits) |  |  | Local scale<br>(*N* = 583 insect species, 18,683 individuals, 51,855 flower visits) |  |  |
| --- | --- | --- | --- | --- | --- | --- |
|  |  |  | % |  |  | % |
|  | %<br>Species | %<br>Individuals | Flower<br>visits | %<br>Species | %<br>Individuals | Flower<br>visits |
| Hymenoptera | 41.30 | 47.65 | 61.90 | 41.51 | 40.36 | 53.47 |
| Diptera | 27.81 | 20.74 | 19.13 | 27.27 | 25.49 | 22.40 |
| Coleoptera | 15.15 | 17.78 | 6.83 | 15.27 | 19.84 | 8.72 |
| Lepidoptera | 12.78 | 13.14 | 11.89 | 13.55 | 13.71 | 15.15 |
| Orthoptera | 1.42 | 0.40 | 0.14 | 0.69 | 0.08 | 0.03 |
| Hemiptera | 1.30 | 0.25 | 0.09 | 1.37 | 0.42 | 0.18 |
| Neuroptera | 0.12 | 0.04 | 0.02 | 0.17 | 0.09 | 0.04 |
| Dermaptera | 0.12 | 0.002 | 0.001 | 0.17 | 0.01 | 0.002 |

Notes: Data for all plant species and dates are combined within each spatial scale. Those for the regional scale are from 288 plant species and 56 sites, and the local scale from 93 plant species at the NC site.

### Pollinator abundance metrics

Abundance ranks of pollinator species for all sampling dates, sites and plant species combined were computed separately from the total number of individuals and flowers visited by each species (see Herrera 2026 for abundance ranks).

Ranks based on individuals and flowers visited were closely correlated at both the regional (*r* = 0.951, *N* = 845, *P* < 2.2E-16) and local spatial scales (*r* = 0.935, *N* = 583, *P* < 2.2E-16). The fitted linear regressions were almost indistinguishable from *y* = *x* isolines (Appendix 1: Figure 1), showing that in practice the two metrics of pollinator abundance were interchangeable. Only pollinator abundances based on number of individuals will be considered hereafter.

**Figure 1.**
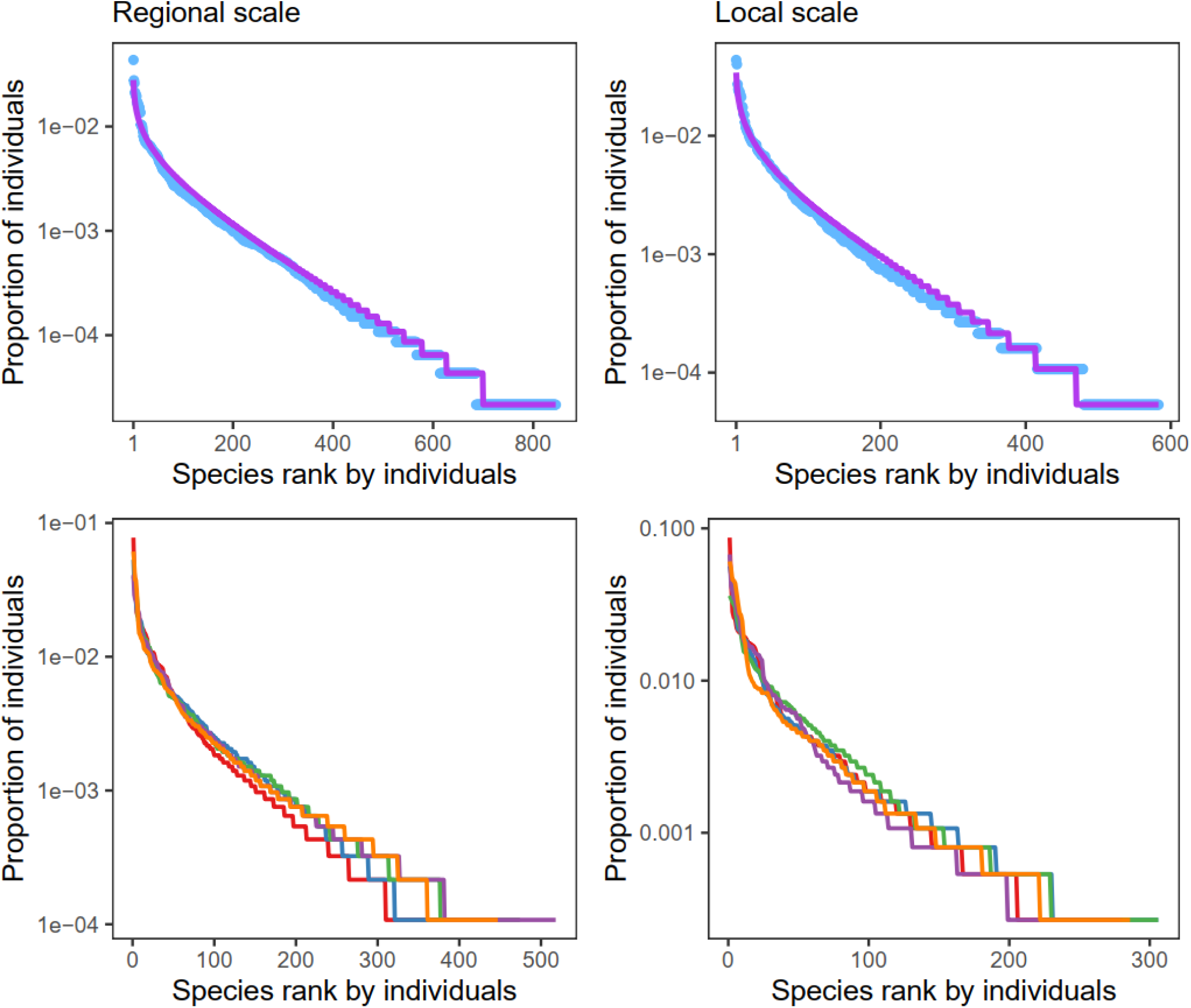
Abundance-rank relationships ("Whittaker plots") for the insect pollinator assemblages at the regional (left) and local (right) spatial scales. Top row: Whittaker plots for the entire sample of pollinators, all plants, sites and dates combined (blue dots). Purple lines depict the predicted relationships from the log-series, the distribution which best fitted the data (Table 2). Bottom row: Differently-colored lines in each graph are separate Whittaker plots for data subsets obtained by splitting the study period into five sequential subperiods having equal numbers of pollinator individuals. Subperiod boundaries were 31 July 1997, 8 July 2012, 31 May 2016, 29 June 2018, 8 June 2022 and 18 October 2025 for regional scale data (*N* = 9,274 pollinator individuals per subperiod), and 26 June 2004, 11 August 2014, 15 July 2016, 10 May 2019, 25 July 2023 and 18 October 2025 for the local scale data (*N* = 3,736 individuals per subperiod).

**Table 2.** Information criteria to compare the performance of five statistical distributions to predict empirical species abundance distributions of insect pollinators. Models are listed in decreasing order of performance. See Figure 2 for abundance-rank diagrams comparing empirical abundance distributions with predictions from the log-series model, and Appendix 1: Table 1 for results of a similar analysis conducted separately for each major insect order.

| Statistical model | Regional scale |  |  |  | Local scale |  |  |  |
| --- | --- | --- | --- | --- | --- | --- | --- | --- |
| | AIC | $\Delta$ | df | w | AIC | $\Delta$ | df | w |
| Log-series | 7245.3 | 0.0 | 1 | 0.550 | 4550.5 | 0.0 | 1 | 0.491 |
| Neutral metacommunity | 7245.7 | 0.4 | 1 | 0.450 | 4550.8 | 0.3 | 1 | 0.421 |
| Poisson lognormal | 7275.3 | 30.0 | 2 | <0.001 | 4553.9 | 3.4 | 2 | 0.088 |
| Power law | 7487.6 | 242.3 | 1 | <0.001 | 4711.0 | 160.5 | 1 | <0.001 |
| Negative binomial | 8895.1 | 1649.9 | 2 | <0.001 | 4936.1 | 385.7 | 2 | <0.001 |

Notes: Data from all plant species and dates combined at regional (56 sites, $N = 845$ pollinator species) and local scales (NC site, $N = 583$ pollinator species). Pollinator abundance was assessed by the number of individuals. Notation for information-theoretical parameters follows Burnham and Anderson (2002). AIC, Akaike's Information Criterion; $\Delta$ , AIC difference, or the increment of AIC relative to the minimum value; df = degrees of freedom, equal to the number of free parameters in each statistical model; w = Akaike's weight, or the relative likelihood of the model given the data, normalized to sum to 1 and interpretable as probabilities.

### Species abundance distributions

The shapes of abundance-rank curves for pollinator species ("Whittaker plots") were virtually identical at the regional and local scales (Figure 1, top row). Ranking species abundance models by AIC differences (Δ_i_) and Akaike weights (*w*_i_) shows that the log-series distribution was the best predictor of empirical pollinator abundance, followed closely by the neutral metacommunity distribution (Table 2). Models based on Poisson lognormal, power law and negative binomial distributions had essentially no empirical support. Results were closely similar when separate analyses were conducted on the species abundance distributions of the four major insect orders (Appendix 1: Table 1).

The most abundant pollinators belonged to a restricted group of species (Figure 1, top row). At regional scale, the five dominant pollinators in decreasing order of abundance were *Bombus terrestris* (Apidae), *Sphaerophoria scripta* (Syrphidae), *Heriades crenulatus* (Megachilidae), *Lasioglossum marginatum* and *Halictus scabiosae* (Halictidae) (Figure 2). These few top pollinators represented only 0.6% of total species but accounted for 13.9% of individuals. On the opposite end of the species abundance gradient were many rare pollinators seen only exceptionally during the study. Out of the 845 insect species comprising the observed pollinator assemblage, 160 (18.9%) were represented by single individuals, and 356 (42.1%) by ≤ 5 individuals. These rare species boosted the overall species richness of the regional pollinator pool, but their contribution in terms of number of individuals was negligible. The set of "right-tail species" was dominated by Hymenoptera (136 species), Diptera (105) and Coleoptera (55), at frequencies roughly similar to those at which they occurred in the pollinator species set at large (Table 1). When separate Whittaker plots were drawn for subsets obtained by splitting data into sequential temporal segments with equal number of pollinator individuals, the form of species abundance distributions remained consistent over the duration of the study (Figure 1, bottom).

**Figure 2.**
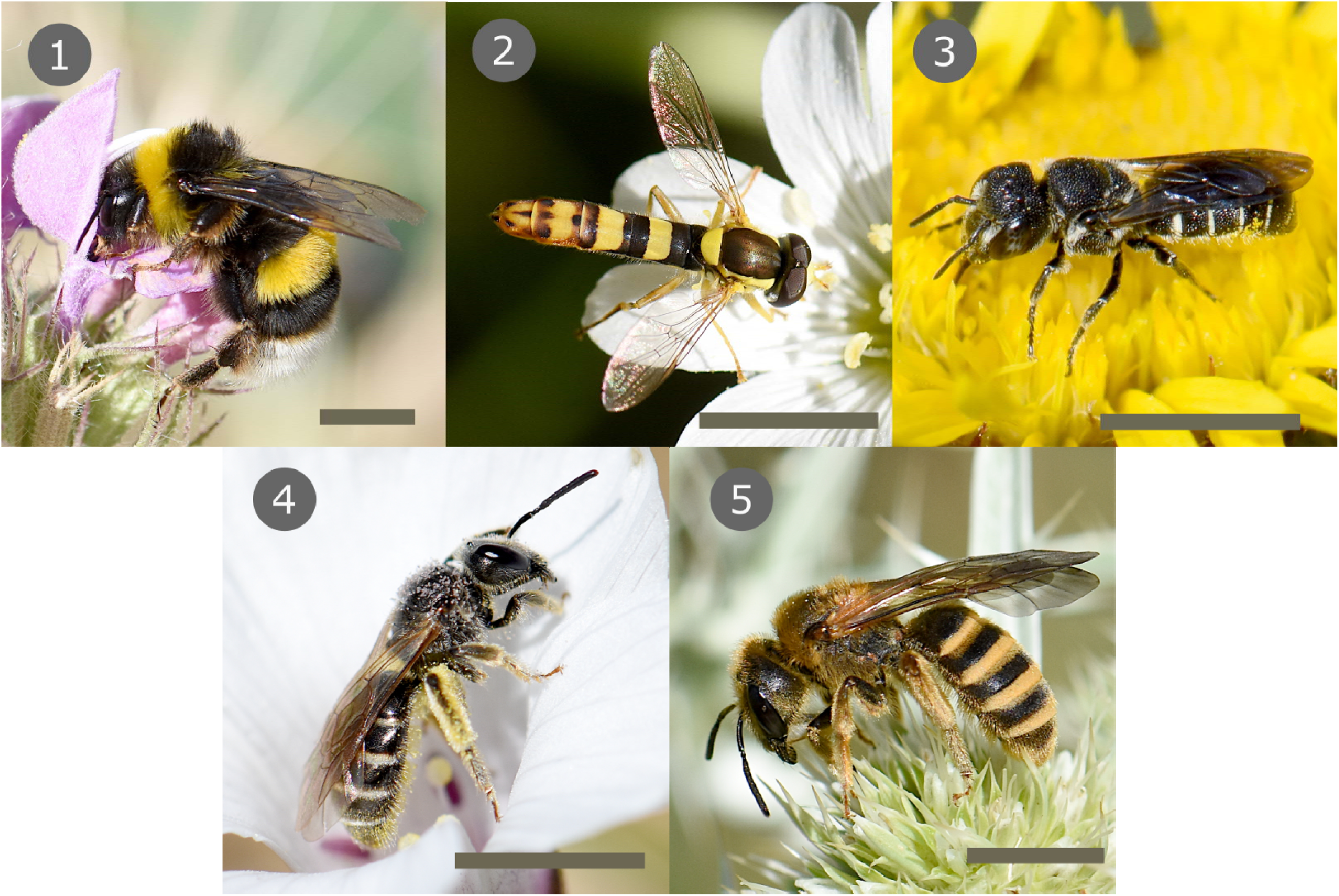
The five most abundant pollinators (data from all plant species combined), collectively representing only 0.6% of all species but accounting for 13.9% of all individuals. 1, *Bombus terrestris* (Apidae); 2, *Sphaerophoria scripta* (Syrphidae); 3, *Heriades crenulatus* (Megachilidae); 4, *Lasioglossum marginatum* (Halictidae); and 5, *Halictus scabiosae* (Halictidae), mentioned in decreasing order of abundance. Scale bar = 5 mm. Photo credit: Carlos M. Herrera.

### Plant intrinsic features and pollinator diversity

#### Sample quality

The first three Hill numbers (Species richness, Shannon diversity and Simpson diversity; ^0^D, ^1^D and ^2^D hereafter, respectively) were estimated for each of the 472 sampling occasions. Sample coverage (SC) was estimated for each sampling occasion as a measure of sampling adequacy. The median and interquartile range of the SC distribution were 0.934 and 0.881-0.963, respectively. These figures indicate that the sampling scheme performed well at capturing pollinator diversity in most sampling occasions, but SC was consistently low in some sampling occasions which yielded very few pollinator individuals due to extremely low pollinator frequency. To reduce biases in ^0^D, ^1^D and ^2^D due to these low-frequency pollinator samples, only the *N* = 447 sampling occasions with SC ≥ 0.75, which comprised data for *N* = 275 plant species, will be included in diversity analyses hereafter. There were weak inverse relationships between SC and diversity measures across sampling occasions in this dataset (Appendix 1: Figure 2), but variation in SC accounted for negligible proportions of variance in diversity (*R*^2^ = 0.019, 0.057 and 0.062, for ^0^D, ^1^D and ^2^D, respectively).

### The pollinator diversity continuum

Plant species means for ^0^D, ^1^D and and ^2^D were obtained by averaging over sampling occasions. Frequency distributions are shown in Figure 3, and summary statistics in Appendix 1: Table 2. Pollinator diversity ranged widely among plant species: Species richness between 1-62 species, Shannon diversity between 1-31 effective species, and Simpson diversity between 1-20 effective species (for concision, I will refer hereafter to units of Shannon and Simpson diversities just as "species"). Hartigans’ dip tests failed to reject the null hypothesis of unimodality for any of three distributions (Dip statistic = 0.025, 0.022 and 0.016, *P* = 0.25, 0.47 and 0.93, for ^0^D, ^1^D and ^2^D, respectively), which is consistent with the impression conveyed by Figure 3 that variation among plant species in pollinator diversity was essentially continuous and unimodal. The three diversity measures were similar in having right-skewed distributions. Most plant species fell near the lower end of the pollinator diversity gradients, and only a minority had high pollinator diversity. For instance, 109 plant species (40% of total) had each ≤10 observed pollinator species, while only 25 plant species (9% of total) had ≥30 pollinator species (Figure 3).

**Figure 3.**
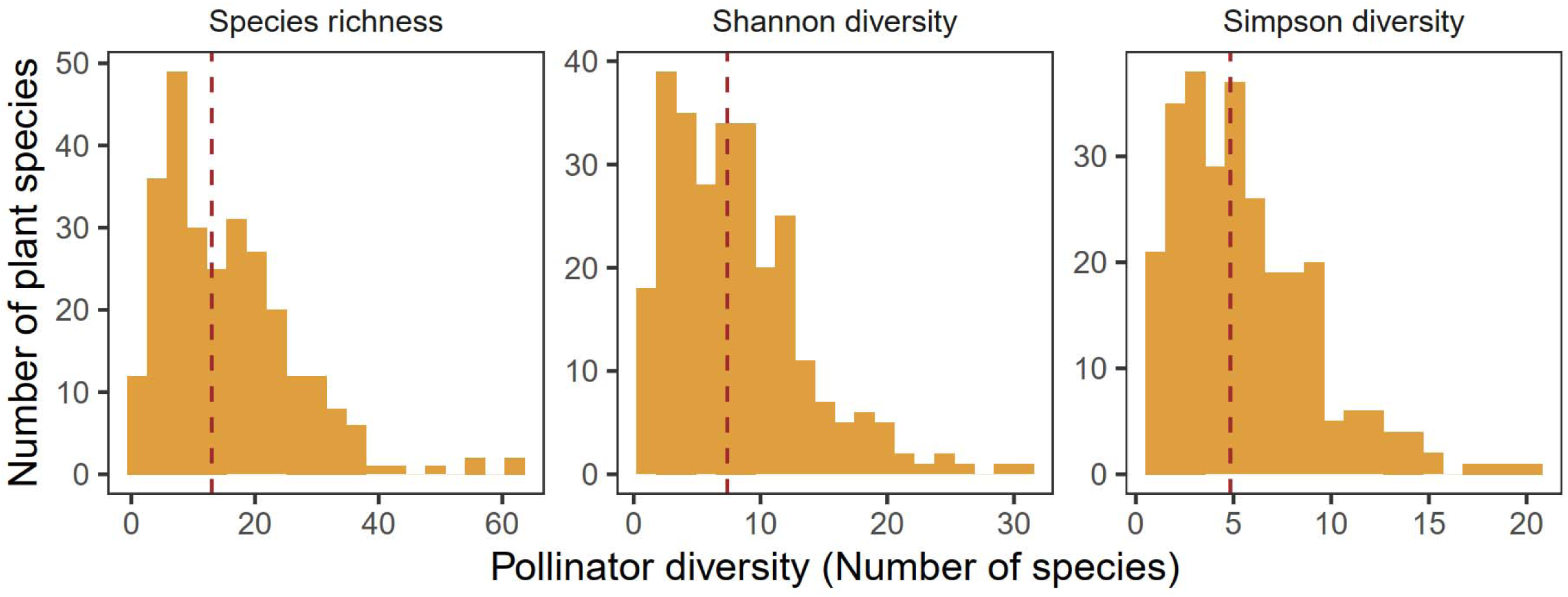
Frequency distributions of Species richness, Shannon diversity and Simpson diversity (Hill numbers ^0^D, ^1^D and ^2^D, respectively) for the pollinator assemblages of *N* = 275 plant species. Vertical dashed lines indicate the medians of distributions.

### Pollinator diversity and plant phylogeny

Variation among plant species in the three measures of diversity exhibited strong, highly significant phylogenetic signals (Pagel’s λ = 0.788, 0.642 and 0.522 for ^0^D, ^1^D and ^2^D, respectively; *P* < E-05 in all cases). This becomes apparent when diversity measures are mapped on the plants’ phylogenetic tree. All pollinator diversity measures were consistently above the medians of their respective distributions in certain plant clades characterized by highly diverse pollinator assemblages (e.g., Asteraceae, Convolvulaceae, Rosaceae), while the reverse held true for other particular clades (e.g., Fabaceae, Brassicaceae, Plantaginaceae) (Figure 5, Appendix 1: Figure 3). Partition of sample-wide variance in pollinator species richness among nested taxonomic categories revealed that variation among plant families and genera within families accounted for 32.5% and 26.9% of ^0^D variance, 22.3% and 23.5% of ^1^D variance, and 16.0% and 20.5% of ^2^D variance, respectively. These figures further support a deep phylogenetic structure in the variation among plant species of pollinator diversity.

### Pollinator diversity and floral features

Linear mixed-effects models fitted to plant species means of pollinator diversity measures, with floral features and their interaction as fixed-effect predictors and plant family as a random effect to control for family-level phylogenetic signal, consistently revealed statistically significant effects on pollinator diversity of perianth type and type of visitation unit, but not their interaction (Appendix 1: Table 3). On average, plant species with flowers aggregated into packets, or those with open perianths, tended to have more diverse pollinator assemblages than those whose visitation units were single flowers or had restrictive perianths (Figure 6, Appendix 1: Figure 4).

**Figure 4.**
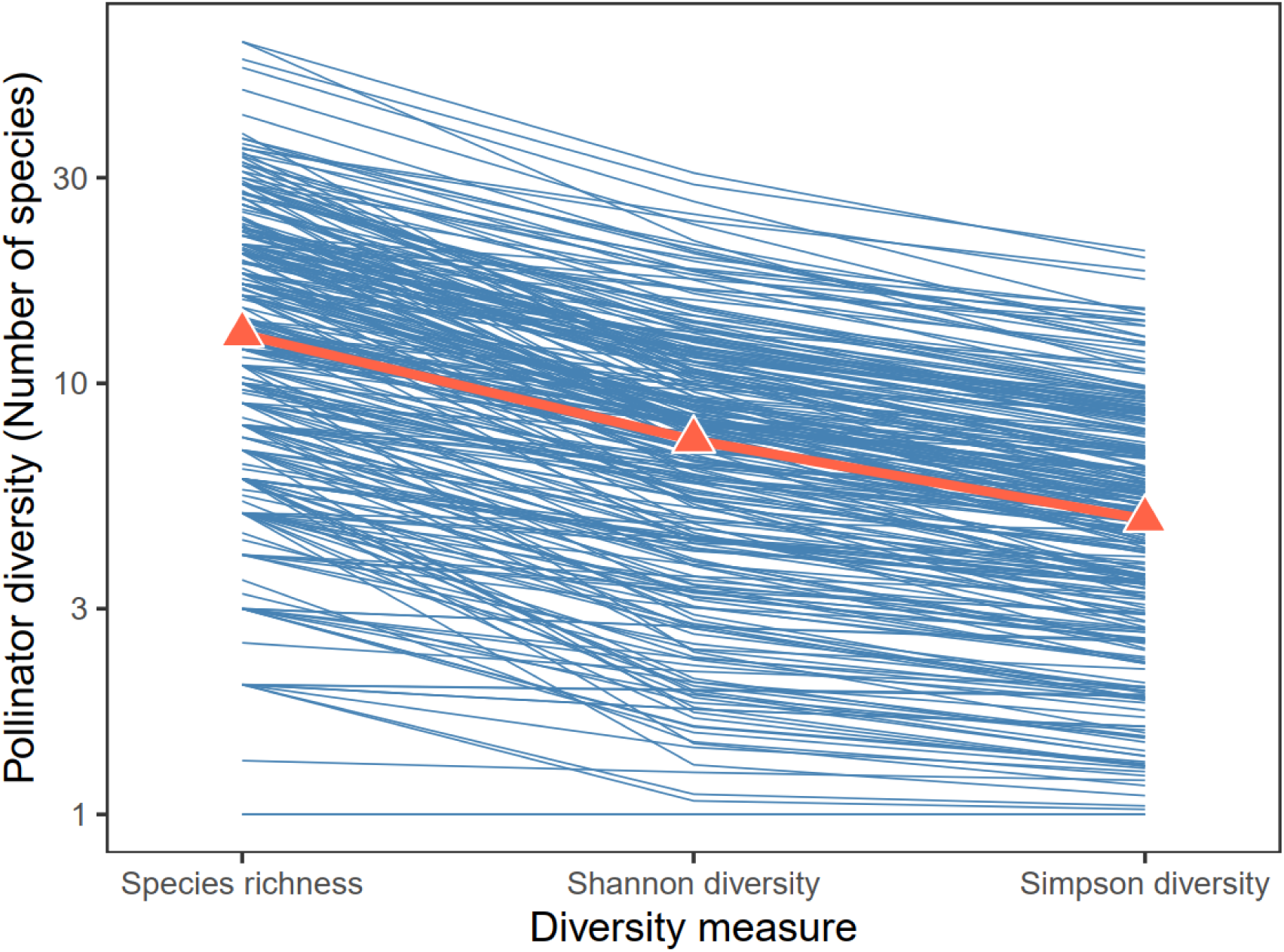
Variation among plant species (*N* = 275) in the relationship between the three measures of pollinator diversity (Species richness, ^0^D; Shannon diversity, ^1^D; Simpson diversity, ^2^D). Each thin blue line corresponds to a different plant species and triangles are the medians of diversity measures. Note logarithmic scale on vertical axis.

**Figure 5.**
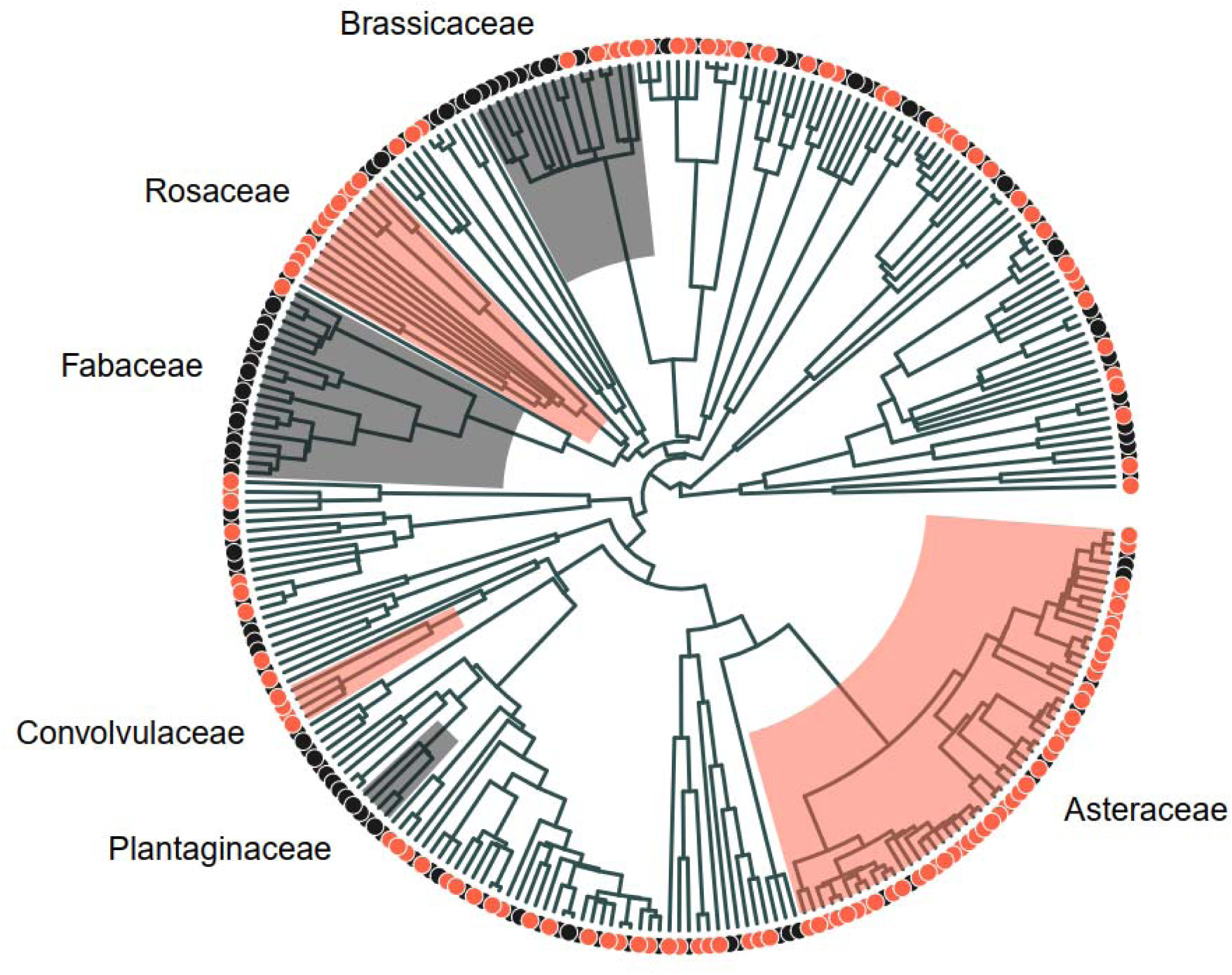
Phylogenetic relationships of the *N* = 275 plant species included in the diversity analyses, and variation across plant species in pollinator species richness. Tree tips are color-coded to denote whether the pollinator species richness of the plant species involved falls above or below the median of the overall distribution (red and grey dots, respectively). Selected clades exemplifying predominantly high (Asteraceae, Convolvulaceae, Rosaceae) or low (Brassicaceae, Fabaceae, Plantaginaceae) pollinator diversity are highlighted (red and grey sectors, respectively). See Appendix 1: Figure 3 for similar graphs corresponding to Shannon and Simpson diversities.

**Figure 6.**
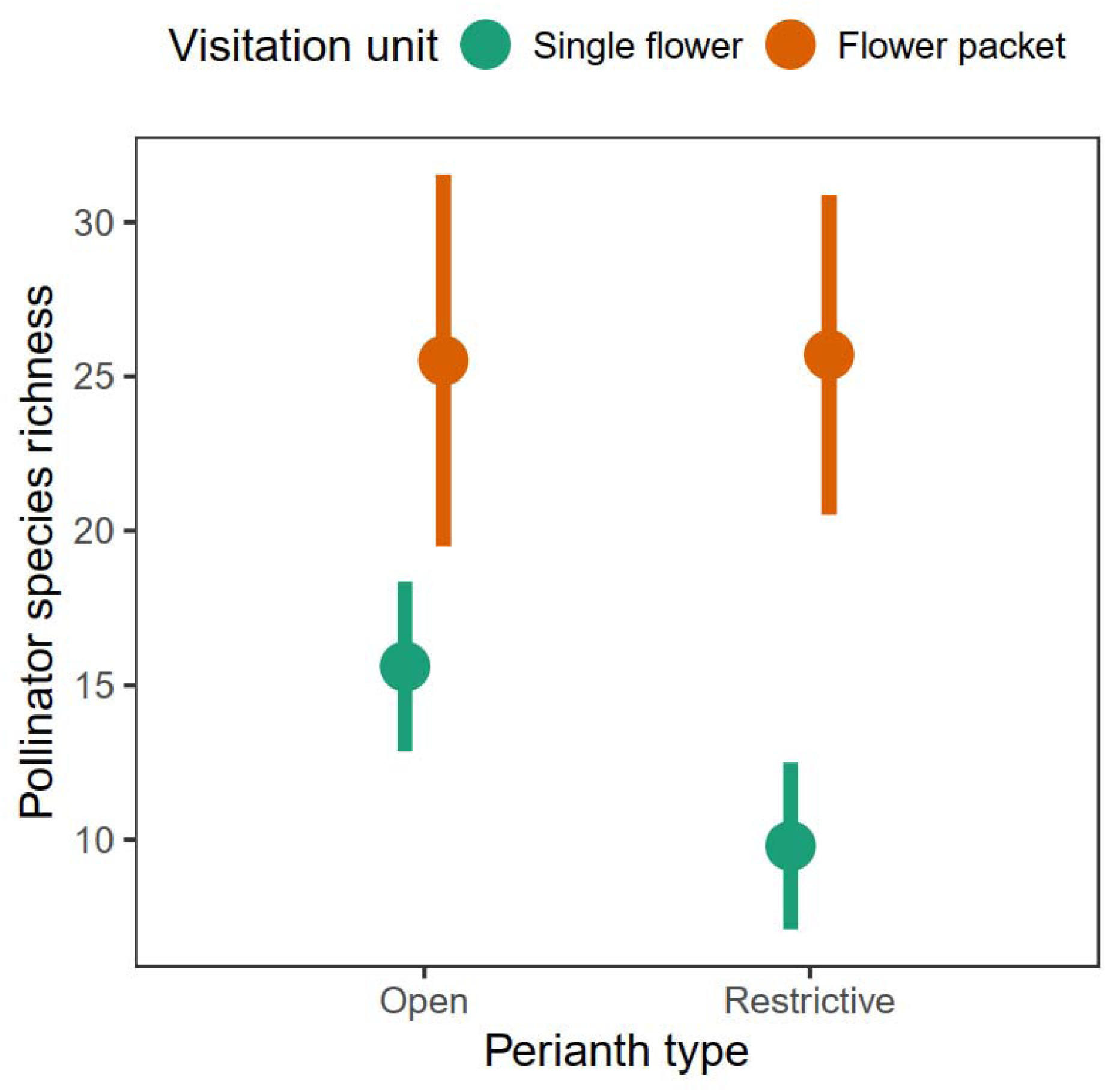
Interaction plot showing model-adjusted, estimated marginal means (dots, vertical segments are 95% confidence intervals) of pollinator species richness for plant species differing in perianth type (open *vs.* restrictive) and pollinator visitation unit (single flower *vs.* flower packet). See Appendix 1: Figure 4 for the graphs corresponding to Shannon and Simpson diversities, and Appendix 1: Table 3 for analytical results.

### Ecological correlates of pollinator diversity

*Pollinator diversity and flower visitation.–* The three measures of pollinator diversity (^0^D, ^1^D and ^2^D) were correlated with the probability per time unit of flowering patches or individual flowers being visited by pollinators (Appendix 1: Table 4). There were consistently nonlinear, convex increases in patch and flower visitation probabilities (all pollinators combined) with increases in pollinator diversity (Figure 7). Relationships depicted are not meant, however, to imply a particular direction of pairwise causality but just to depict the shape of bivariate relationships between variables. Pollinator diversity was more closely correlated with patch (deviance = 36-75%) than flower visitation (deviance = 14-19%) probability (Appendix 1: Table 4).

**Figure 7.**
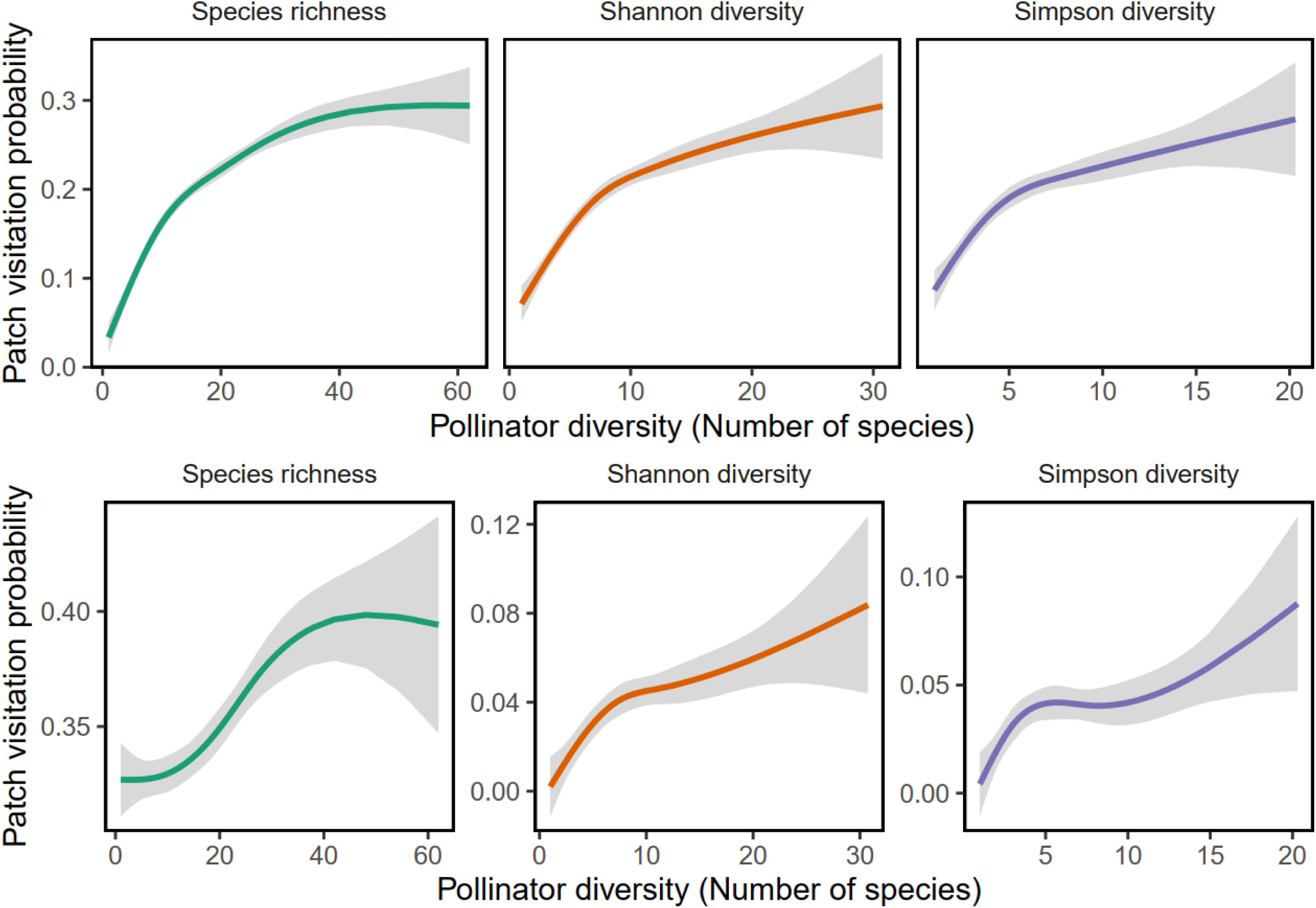
Relationships across plant species between two measures of pollinator visitation rate and three measures of pollinator diversity, obtained by fitting nonparametric regressions to species means using smoothing splines. Pollinator visitation rates are per-min probability estimates of some pollinator visiting one flowering patch (top row) or one individual flower (bottom row). Lines are the estimated marginal effects of diversity on visitation rates after statistically accounting for interspecific variation in sample coverage (Appendix 1: Figure2), and shaded bands are 95% confidence envelopes around the marginal effect estimate. See Appendix 1: Table 4 for analytical details.

### Pollinator diversity and habitat type

Linear mixed-effects models were separately fitted to plant species means for ^0^D, ^1^D and ^2^D, with habitat type as the single fixed-effect predictor, and plant family and genus-nested-within family as random effects to control for phylogenetic effects. Every measure of pollinator diversity varied significantly among habitat types after accounting for differences among habitats in taxonomic composition of plant communities (Appendix 1: Table 5). Estimated marginal means of pollinator diversity were highest in forest clearings and edges, and lowest in dolomitic outcrops, rock cliffs and forest interior (Figure 8). Variation among habitats in pollinator diversity involved changes in the relative values of the three diversity measures. Pollinator species richness was much higher than Shannon and Simpson diversities in all habitats except dolomitic outcrops, rock cliffs and forest interior, where the three diversity measures fell within rather narrow limits (Figure 8).

**Figure 8.**
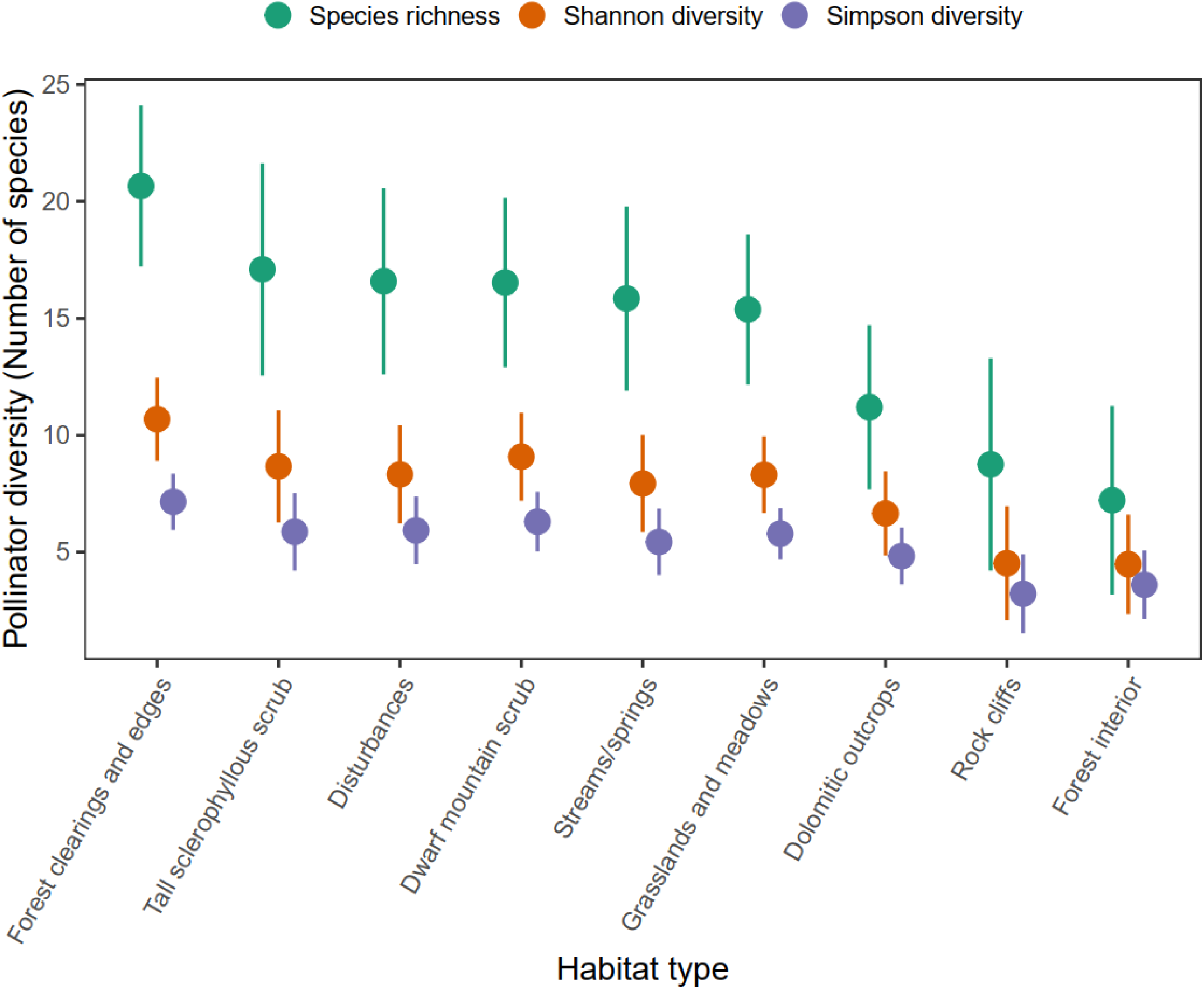
Model-adjusted, estimated marginal means (dots, vertical segments are 95% confidence intervals) of pollinator diversity in the nine structural habitat types recognized in this study. Linear mixed-effects models were fitted testing for the effect of habitat type on the plant species means for the three pollinator diversity measures while statistically accounting for differences among habitats in taxonomic composition of plant communities. Habitat types are arranged from left to right in decreasing order of pollinator diversity. See Appendix 1: Table 5 for analytical details.

#### Temporal dimension of pollinator diversity

Generalized additive mixed models were fitted to the pollinator data from the *N* = 447 sampling occasions to test for seasonal and supra-annual changes in pollinator diversity. Smoothers of sampling year and time of year were the models’ predictors. Plant species and sampling site were included as random effects to account for statistical non-independence of diversity data for the same plant species or sampling site. Analytical results of models are summarized in Appendix 1: Table 6.

Pollinator diversity exhibited a distinct seasonal rhythm whose shape was roughly the same for all diversity measures, as denoted by the close similarity of estimated degrees of freedom of smoothing terms (Appendix 1: Table 6). On average, diversity increased slowly from the beginning of year up to around day 150 (late May), then increased faster until reaching an annual peak around day 225 (late July-early August), and steadily declined afterwards until reaching minimal values by the end of year (Figure 9). The fast building-up of pollinator diversity which took place in late spring-early summer was proportionally stronger for species richness than for the other diversity measures (Figure 9), implying a higher dominance during that period.

**Figure 9.**
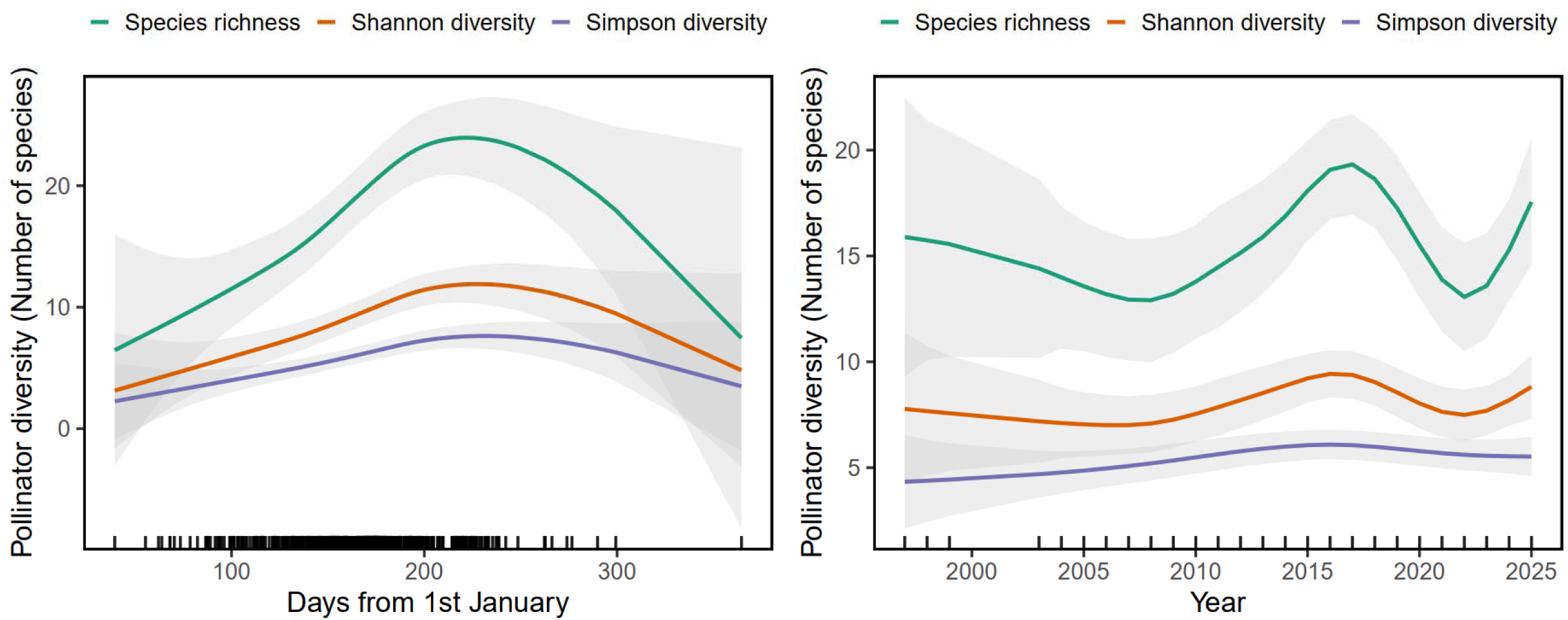
Seasonal (left) and interannual (right) trends in mean predicted pollinator diversity at the plant community level. Lines are smoothers obtained from generalized additive mixed-effects models fitted to pollinator diversity measures from the *N* = 447 plant species x site x year combinations represented in the data set, and shaded bands are 95% confidence envelopes around the prediction. See Appendix 1: Table 6 for analytical details of generalized additive mixed models.

Species richness and, to a lesser extent, Shannon diversity of the pollinator assemblage fluctuated over the 1997-2025 study period (Appendix 1: Table 6), showing two distinct dips (2008 and 2022) and one peak (2017) (Figure 9). Model-adjusted, mean estimated species richness for the plant community as a whole ranged between 12.9 and 19.2 species in the lowest- and highest-diversity years (2008 and 2017, respectively), or about 50% variation relative to the lowermost, 2008 figure. Simpson diversity, in contrast, remained fairly constant around ∼5 species throughout the study (Figure 9). The contrasting shapes of long-term fluctuations in the three diversity measures is analytically supported by the steep decline in estimated degrees of freedom of the smoothers for year in the direction Species richness–Shannon diversity–Simpson diversity (Appendix 1: Table 6).

### Appraising "dark diversity"

Predicted accumulation curves of pollinator diversity with increasing number of individuals observed are illustrated in Figure 10 for all taxa combined and for each major insect order. All curves for a given diversity measure looked virtually identical when axes were adjusted for differences in scale. Shannon and Simpson diversity measures consistently reached a plateau at small numbers of sampled individuals. Pollinator species richness, in contrast, did not reach a plateau, hence suggesting that a non-trivial proportion of existing species of all major insect orders remained undetected despite the large sample sizes of this study. Theoretical estimates of the proportion of total existing species which remained undetected, computed as (Predicted richness - Observed richness)/Predicted richness are summarized in Table 3. Although the two estimation procedures produced slightly different numerical results, the overall picture was similar: on average, ∼16% of existing species were missed. Dark diversity was highest for Diptera and Coleoptera, and lowest for Hymenoptera and Lepidoptera (Table 3). For all taxa combined, the predicted number of total insect pollinator species in the area studied was ∼1,000 species.

**Figure 10.**
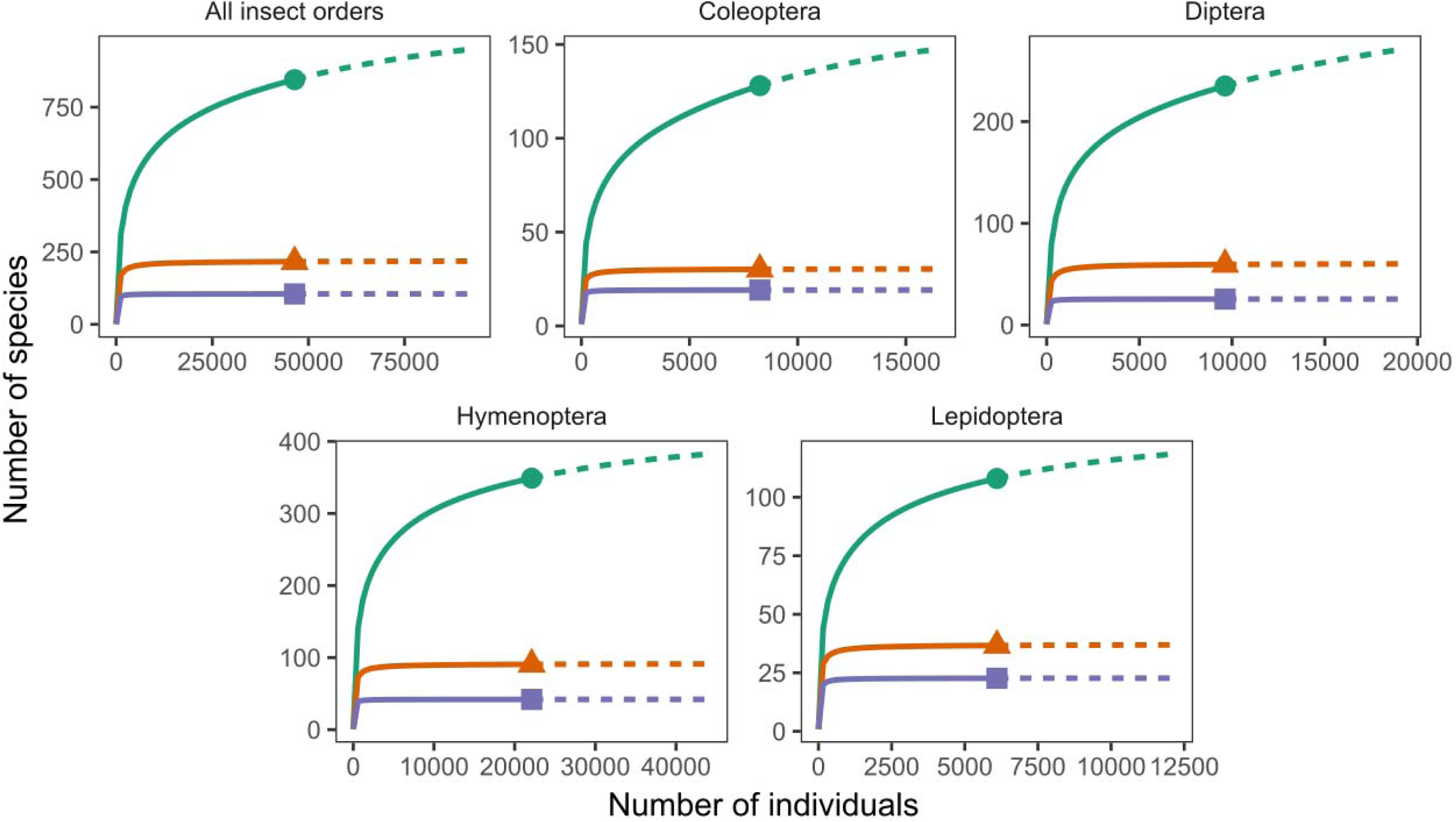
Curves of cumulative pollinator diversity with increasing number of individuals for the entire data set and the four most abundant insect orders, obtained by interpolation-extrapolation. Note that the scales of horizontal and vertical axes differ among graphs. Symbols (dots, triangles and squares, for Species richness, Shannon diversity and Simpson diversity, respectively) indicate the observed, empirical diversity values. Continuous and dashed lines correspond to the rarefaction and extrapolation segments of curves, respectively.

**Table 3.** Observed and predicted pollinator species richness, and estimated proportion of undetected species ("dark diversity") for the four major insect orders and the complete data set.

| Insect group | Species<br>observed | Estimation method |  |  |  |
| --- | --- | --- | --- | --- | --- |
|  |  | Interpolation-extrapolation |  | Good-Turing |  |
|  |  | Predicted<br>± SE | %<br>Undetected | Predicted<br>± SE | %<br>Undetected |
| Hymenoptera | 349 | 401 ± 18 | 13.0 | 403 ± 11 | 13.4 |
| Diptera | 235 | 308 ± 19 | 23.8 | 285 ± 11 | 17.5 |
| Coleoptera | 128 | 160 ± 15 | 20.1 | 158 ± 8 | 19.0 |
| Lepidoptera | 108 | 124 ± 11 | 12.9 | 125 ± 5 | 13.6 |
| All data | 845 | 1,023 ± 32 | 17.4 | 1,005 ± 19 | 15.9 |

*Notes:* The predicted number of existing species was obtained using the interpolation-extrapolation method of Chao et al. (2014) as implemented by Hsieh et al. (2025) in the R package iNEXT, and the Good-Turing method of Chao et al. (2017) as implemented in the online SuperDuplicates application (https://chao.shinyapps.io/SuperDuplicates/; accessed December 2025). The estimated proportion of total existing species that remained undetected during this study was computed as (Predicted richness - Observed richness)/Predicted richness.

## DISCUSSION

### Challenges to pollinator community research

The set of pollinator species that coexist sympatrically can be treated as an ecological community despite their taxonomic heterogeneity because they are all consumers of the same trophic resources (nectar and pollen). The concepts and tools of community ecology traditionally applied to other biological communities could therefore be applied to pollinators, as exemplified by this and other studies (Tepedino & Stanton 1981; Kalin Arroyo et al. 1982; Kato et al. 1990; Kevan et al. 1997; Norfolk 2015; Roswell et al. 2023). The fact that this has been done so infrequently must be attributed to practical rather than conceptual hindrances.

Although the tradition of surveying local or regional pollinator assemblages dates back to more than one century (Müller 1883; Knuth 1908; Robertson 1928), suitable quantitative data for community analyses are hard to obtain. Cost-effective, unattended sampling methods tend to produce biased compositional data (Hall 2018; Thompson et al. 2021; Moldoveanu et al. 2025; Serra-Marin et al. 2025; Watazu et al. 2025) and usually lack the critical information on the actual pollinating role of trapped species. This leaves watching of pollinator activity at flowers as the method of choice for obtaining unbiased assessments of the species composition of pollinator communities (Herrera 2019; but see Serra-Marin et al. 2025 for the possibility of observer-induced biases). Apart from being time-consuming and cost-ineffective, this procedure is problematic in very species-rich plant communities. To be reliable, pollinator composition data at the plant community level should be obtained for the majority of animal-pollinated plants, not just arbitrarily defined, small species subsets prone to biases due to taxonomic affiliation or floral features (Ollerton et al. 2015; Ollerton 2017; Herrera 2020; Adamo et al. 2021). For example, in the extensive set of European plant-pollinator networks (*N* = 1,522) assembled by Lanuza et al. (2025), the median numbers of plant and pollinator species per network were only 9 and 12 species, respectively (interquartile ranges = 5-15 and 7-24 species for plants and pollinators, respectively), which denotes the generally feeble coverage of local or regional plant and pollinator diversities in most of these "community" studies.

Reliable identification of pollinators to species is also central to any pollinator diversity analysis, and here too automated methods fall short at producing "species grade" pollinator data (Serra-Marin et al. 2025; Watazu et al. 2025). Accurate, thorough species-level identification of pollinators is most challenging when large numbers of species from widely disparate taxonomic groups are involved, as exemplified by the 845 species found in this study, the 715 species in a temperate beech forest in Japan (Kato et al. 1990), or the 666 species from an eastern Mediterranean phryganic ecosystem (Petanidou 1991; Petanidou & Ellis 1993). The problem of species identification can be almost insuperable in hyperdiverse regions with limited taxonomic resources (Herrera et al. 2025). Grouping undetermined numbers of unnamed species under arbitrarily defined umbrellas such as "morphospecies" or "pollinator functional groups" has been sometimes used to overcome species identification problems. This can be a sound procedure for some purposes, but it precludes valid analyses of species abundance distributions and diversity. By combining long-term intensive fieldwork under the ecology-of-place research paradigm (Price & Billick 2010) with thorough pollinator identification to species, this study was able to address the seemingly simple but still largely unformulated questions outlined in the Introduction.

### Species abundance distributions

Species abundance distributions (SADs) provide quantitative representations of the full distribution of commoness and rarity in multispecies assemblages, which explains why SADs are one of the most widely studied and reviewed patterns in ecology (Hubbell 2001; McGill et al. 2007; Alroy 2015; Baldridge et al. 2016; Su 2018). The rarity of pollinator communities as focal subjects of SAD studies stressed in the Introduction is most likely a consequence of the real-world problems outlined in the preceding section. I am only aware of a few exceptions.

Tepedino and Stanton (1981) studied dominance-diversity curves for bees in a short-grass prairie and found that they fitted a log-normal distribution. Pollinator species abundances did not depart from log-normal distributions in the small sample of pollination studies reviewed by Herrera (1989). Kevan et al. (1997) showed that bees from pesticide-free blueberry fields fitted well the log-normal model of species diversity and abundance, but not those from fields surrounded by pesticide-treated forest. Kato et al. (1990) explored dominance-diversity curves of all insect pollinators, not just bees, from a beech forest but their study was not a full-fledged SAD analysis because infrequent species, which can be more important than frequent ones as determinants of SAD patterns (Gotelli & Chao 2013; Chao et al. 2017), were left unaccounted. The present study thus seems the first comprehensive attempt to date at elucidating SAD for a throughly sampled, taxonomically comprehensive insect pollinator community.

The empirical SAD of the insect pollinator assemblage in the Sierra de Cazorla did not depart significantly from those of other multispecific insect assemblages defined by taxonomy (order, family) or geographic location (region, sampling site). The numerical prevalence of a few dominant species and the profusion of rare species found here is a long-known, ubiquitous pattern exhibited by any sufficiently-sampled insect assemblage (Corbet 1941; Fisher et al. 1943; Williams 1964; Kempton & Taylor 1974). Certain similarities between results of this study for insect pollinators and those reported elsewhere for other large insect samples are remarkable. For example, the proportion of total pollinator species represented by singletons was 19% in this study, the same as for tropical butterflies from the Malay Peninsula (19%, Corbet 1941) or moths captured in an English light trap (19%, Williams 1964: Table 8). The corresponding figures for the proportion of species represented by ≤5 individuals were 42%, 47% and 48%, respectively, again closely similar.

Initially derived as a purely statistical distribution by Fisher et al. (1943), the log-series model of species diversity and abundance has been subsequently related to ecological processes, particularly in relation to the neutral theory of ecological diversity, which emphasizes stochastic processes in structuring ecological communities and challenges traditional niche-based views of ecology (Hubbell 2001; Alonso & McKane 2004; Matthews & Whittaker 2015). As frequently found in previous research on SADs of insect assemblages (Ulrich et al. 2010; Baldridge et al. 2016), the log-series was also here the statistical model that best predicted pollinator species abundances at local and regional scales. If the log-series is actually the predominant statistical model for species diversity and abundance in large multispecific collections of insects, then subsamples drawn from the "insect sampling universe" comprising only species which behave as pollinators should also fit a log-series for no particular reason other than passive "inheritance" of statistical properties from the sampling universe (Williams 1964). May (1975, p. 95) likewise noted that "[the logseries distribution] has the elegant property that samples taken from a population distributed according to a logseries are themselves logseries". This property stems from the fact that under a log-series distribution, species abundances in the community are independent identically distributed log-series variables (Prado et al. 2025). If the SADs of terrestrial insect communities as a whole generally conform to the the log-series model, then the SADs of insect pollinator assemblages are expected to conform to the log-series distribution just as a by-product of the distribution’s "elegant property". Strong support to this view is provided here by the close similarity of the results of SADs analyses conducted independently on the four major orders of insect pollinators (Appendix 1: Table 1).

Although the log-series was the model which consistently provided the best prediction of the SADs examined in this study, the close similarity of Akaike weights for the log-series and the neutral metacommunity distribution (Table 2) indicates that the best model is not convincingly best (Burnham & Anderson 2002). The possibility cannot be dismissed therefore that the empirical SAD for pollinators actually conforms best to expectations explicitly derived from the neutral theory of ecological diversity assumed by the neutral metacommunity distribution (Hubbell 2001; Alonso & McKane 2004). The difficulty for discriminating between the predictive performance of the two best models in Table 2 most likely reflects that the neutral metacommunity distribution converges to the log-series when diversities are as high as in the pollinator data set analyzed here (Alonso & McKane 2004).

### Pollinator diversity: plant community patterns

A purported dichotomy involving "generalized" *vs.* "specialized" pollination is frequently alluded to in studies of plant-pollinator systems when comparing species or populations with many *vs.* few species of pollinators (Herrera 2005; Waser & Ollerton 2006; Gómez et al. 2007, 2015; Valverde et al. 2019). The study of an entire community of insect-pollinated plants presented here confirms the reality of broad interspecific variation in pollinator diversity among sympatric plants. At the same time, however, the continuous, unimodal distributions of diversity measures tends to negate the existence of recognizable, discrete groups of plant species with contrasting levels of pollinator diversity. Shoehorning the continuous, unimodal variation of pollinator diversity into informal verbal categories such as "generalist" and "specialist" can hinder a deeper understanding of how plant-pollinator interactions are organized at the community level.

This study has shown that, given sufficient ecological and phylogenetic scope, a substantial part of interspecific variation in the pollinator diversity of a plant community could be parsimoniously predicted by the simultaneous consideration of a few intrinsic plant features and environmentally-defined drivers. On one side, the strong phylogenetic signal and the partition of variance of pollinator diversity into hierarchically nested taxonomic levels indicate that pollinator diversity of a given plant can be largely predicted by its evolutionary history alone. Also supporting a role of intrinsic, species-specific determinants of pollinator diversity is the finding that simple floral features, such as perianth restrictiveness and flower clustering into aggregated structures can predict pollinator diversity. These results are not unexpected, as effects of corolla restrictiveness and floral arrangement on diversity and abundance of pollinators have been frequently documented (Gong & Huang 2009; Montgomery & Rathcke 2012; Blanco-Pastor et al. 2015; Herrera 2020). On the other side, ecological settings as defined by habitat type were good predictors of mean pollinator diversity associated with a given plant. After statistically accounting for variation among habitats in plant taxonomical composition, species living in dolomitic outcrops, rock cliffs and forest interior stood out by the low average diversity of their pollinator assemblages.

These three habitats also had the lowest pollinator abundance (Herrera 2020), hence the same ecological factors were probably responsible for the parallel variation in pollinator abundance and diversity. Low soil fertility and water availability in cliffs and dolomitic outcrops probably constrain the amount and quality of floral rewards produced by plants (Waser & Price 2016; Gallagher & Campbell 2017). The low solar irradiance reaching cliffs and forest understory will probably constrain plant photosynthesis and the production of floral rewards (Lüttge 2013), and also limit foraging opportunities by ectothermic insect pollinators whose activity requires exposure to sunshine (Herrera 2020, 2024).

Plant species with more diverse pollinator assemblages tended to have higher pollinator visitation rates to both flowering patches and individual flowers. It is unlikely that this relationship is a spurious effect of links between diversity and sampling coverage (SC), both because the diversity-SC relationship was very weak and the possible correlation SC-abundance was statistically accounted for by including SC as a covariable in models. Instead, the diversity-visitation correlations most likely arose because, everything else being equal, more pollinator species will mean more individuals and hence greater flower visitation. The convex, nonlinear form of abundance-diversity relationships suggests some underlying "diminishing returns" process, whereby the proportional contribution of increased pollinator diversity to visitation rates declined with increasing diversity, due perhaps to increasing exploitative or interference competition at flowers. These results also have some practical implications. It has been recently proposed (Marini et al. 2025) that the easier-to-estimate total pollinator abundance/frequency can provide a valuable proxy for the harder-to-estimate pollinator diversity. While results shown here clearly support that view, nonlinearity of the abundance-diversity relations should be taken into consideration when making projections of pollinator diversity based on abundance alone. Furthermore, only studies relying on experimental manipulations of pollinator species richness or abundance will be apt to dissect the effects of these two correlated variables on community-level pollinator services or pollination success of plant species or individuals (Albrecht et al. 2012; Fründ et al. 2013; Genung et al. 2017; Artamendi et al. 2025). When such manipulative experiments are impractical, however, the use of ^2^D to measure pollinator diversity would help to alleviate the statistical nuisance introduced by the diversity-abundance relationship, as this is the diversity measure with the weakest relationship with pollinator abundance (Appendix 1: Table 4). As noted by one reviewer, this would be achieved by redefining diversity as something much closer to evenness, thus implying a conceptual shift as well (Mark Genung, *personal communication*).

Spatial and temporal patterns of pollinator diversity documented in this paper are expected to have functional consequences in terms of pollination success and reproductive performance of plants (Artamendi et al. 2025). First, the relationship discussed above between pollinator diversity and patch/flower visitation probabilities provides a straightforward mechanism whereby variations in pollinator diversity will translate into variations in pollen transfer rates and pollination service. This relationship should be modulated by differences among species of pollinators in per-visit probability of pollen pick-up and deposition (Herrera 1987a; King et al. 2013). And second, pollinator diversity can enhance plant reproductive success through complementarity effects (Herrera 2000; Blüthgen & Klein 2011; Albrecht et al. 2012; Fründ et al. 2013; Martins et al. 2015; Loy & Brosi 2022). While functional complementarity effects have been most convincingly shown in relation to bee diversity, they are expected to be even stronger when taxonomically disparate pollinator groups are involved (Herrera 1987a,b, 2000; Albrecht et al. 2012; Fründ et al. 2013; Venjakob et al. 2016).

### Pollinator diversity by Hill numbers

The compositional complexity of a species assemblage is not expressible as a single number, and standard measures such as diversities (Hill numbers) vary depending on the measures’ emphasis on rare or common species as defined by the order *q* (Hill 1973; Chao & Jost 2015). Multispecies assemblages will therefore be better described by the profile of several diversity measures of order *q* ≥ 0 (Gotelli & Chao 2013; Chao & Jost 2015), and this was the analytical strategy adopted in this paper by concurrently examining Hill numbers for *q* = 0, 1, 2. Results have shown that focusing on pollinator species richness (*q* = 0) will miss some ecologically significant patterns of pollinator variation in time and space, as discussed below.

On the one hand, overimposed on an overall pattern of concordance among the three diversity measures (^0^D, ^1^D and ^2^D) there was broad variation among plant species in slope of the profile defined by ^0^D, ^1^D and ^2^D (Figure 4). This was indicative of interspecific heterogeneity in the relative importance of rare pollinators or, in other words, that the dominance of the few most abundant pollinators varied broadly among plant species, in ways that were largely decoupled from variation in pollinator species richness. This points to the importance of incorporating species dominance as an additional descriptive axis in crude comparisons of pollinator species richness. And on the other hand, and possibly most important, the weak coupling of the variation of ^0^D, ^1^D and ^2^D among habitats, years and time of year revealed a deep, complex spatio-temporal patterning of pollinator species’ dominance/evenness in the pollinator assemblages of individual plant species. Habitats differed in slope of the ^0^D–^1^D–^2^D profiles, which were shallower in the least-diverse pollinator assemblages of plants from dolomitic outcrops, rock cliffs and forest interior than in the rest of habitat types (Figure 8). In the same vein, supra-annual and seasonal variations in the relative values of the ^0^D–^1^D–^2^D triplet (Figure 9) likewise denoted seasonal and long-term temporal changes in the dominance of the most abundant pollinators. In general, therefore, ^1^D and ^2^D were found to be less spatially and temporally variable, or more ecologically robust, than ^0^D. This finding is consistent with the suprannual constancy in the shape of species abundance distributions (SADs) at regional and local scales despite substantial long-term fluctuations in ^0^D, and further stresses the value of combining diversity measures to describe pollinator communities, as emphasized by Chao and Jost (2015) for ecological communities in general. It remains to be investigated (but see Roswell et al. 2023) whether variation in pollinator Hill numbers among plant species or populations has predictable consequences for plant pollination and reproductive success through effects on the strength of functional complementarity effects. It seems plausible to predict, however, that the shallower the ^0^D-^1^D-^2^D relationship holding for a given pollinator assemblage (e.g., forest interior plants in this study), the more apparent should the effects of functional complementarity on plant reproduction become, and vice versa.

Estimated proportions of undetected pollinators varied widely among insect orders, being particularly high for Coleoptera and Diptera in comparison to Hymenoptera and Lepidoptera. Reasons for these taxonomic differences in the estimated incidence of undetected species are unclear at the moment, but I suggest that they must be related to intrinsic differences among groups in detectability and species identifiability in the field. Supporting the latter is the parallel variation across insect orders found here between estimated dark diversity and the proportion of individuals which I was eventually unable to identify to species. This suggests that an unknown proportion of undetected species, particularly among Diptera and Coleoptera, were actually detected but could not be identified to species. I propose to name these detected but unidentified species as "crepuscular diversity", whose magnitude will depend on ease of recognition and current state-of-the-art of taxonomic knowledge.

Regardless of the causes, the wide disparity among major pollinator groups in the estimated proportion of dark (including crepuscular) diversity implies that empirical data describing the composition of diverse pollinator assemblages are prone to be systematically biased against taxonomic groups with inherently high dark and crepuscular diversity, and low identifiability. This notoriously applies to dipterans, which are predicted to account by themselves for ∼42% of all undetected species (computed from figures in Table 3). This stands as yet another argument against the unjustified neglect of dipterans in studies of plant-pollinator interactions (Doyle et al. 2020; Garcia et al. 2022).

Direct species counts were once considered to provide one of "the simplest, most practical, and most objective measures of species richness" (Peet 1974, p. 290). Nowadays, in contrast, species richness is seen as "a problematic index of biodiversity for two reasons related to sampling intensity and the species abundance distribution" (Chao et al. 2014, p. 45), whose "statistical properties […] are much worse than those of any other common diversity measure" (Chao and Jost 2012, p. 2533). Results discussed in this section have shown that, for these same two reasons, species richness alone is also a poor measure of diversity when it comes to multispecies pollinator assemblages, and that caution should be exercised when making comparisons or inferences using pollinator diversity estimates based on that measure alone. More generally, this conclusion agrees with Larsen et al.’s (2018) contention that comprehensive assessments of diversity are necessary when long-term ecological changes at the community level are at stake. This becomes particularly relevant in pollination studies, given the current worldwide decline of pollinator diversity in response to accelerating environmental change (Artamendi et al. 2025). As shown in this study, different Hill numbers can display contrasting long-term changes. The substantial supra-annual fluctuation of pollinator species richness probably reflected variations in importance of transient species (*sensu* Taylor et al. 2018) combined with intense supra-annual turnover in species composition (C. M. Herrera, *unpublished data*), whereas the relative invariance across years of Shannon diversity and, particularly, Simpson diversity reflected strong long-term robustness in the statistical architecture of pollinator SADs (see also Hu et al. 2019). Focusing on changes of species richness alone would have produced a misleading picture of the overall long-term changes in pollinator diversity taking place at the plant community level.

## CONCLUDING REMARKS

Mutualistic plant-pollinator systems arguably represent the epitome of the sort of "feedback systems" that Margalef (1968, p. 19) had in mind when he referred to diversity measures as *"an expression of the possibilities of constructing feedback systems or any sort of links, in a given assemblage of species"*, since they exemplify systems whose persistence depends on the combined diversity of the organisms sitting on the two sides of the interaction. Nevertheless, whereas the question "How many plants are pollinated by animals?" has been addressed on innumerable occasions in the wake of Ollerton et al.’s (2011) influential paper, the symmetrical and intrinsically related one, "How many animals pollinate plants?" went essentially neglected during the same period. While answering the first question is essential to understand the importance of animals in the pollination of flowers, the second one is fundamental to evaluate the ecological and evolutionary dimensions of local or regional assemblages of animal pollinators, which in turn can ultimately determine the resilience of plant-pollinator systems to extinctions. Coordinated answers to both questions are indispensable to comprehend the underpinnings of current losses of biological diversity and ecological functionality, and will be essential for devising ecologically meaningful actions for alleviating or reverting biodiversity loss.

If we accept that the ultimate purpose of measuring diversity is, as it was felt by Hill (1973), to gain an objective means of performing comparisons, then only rigorous pollinator diversity measures adhering to a widely accepted "gold standard" will do the job of evaluating long-term trends in "ecological health" of plant-pollinator systems. For example, in the particular case of the hyperdiverse plant-pollinator system studied here, one pollinator diversity estimator (^0^D) fluctuated substantially in the long run while others (^1^D, ^2^D) remained fairly constant, and none evidenced any sustained, long-term decline over the 29 years covered by this study. On one hand, this admittedly complex temporal pattern of diversity emphasizes the perils of sweeping generalizations and unrealistic simplifications of pollinator declines already pointed out by, among others, Herrera (2019, and references therein). On the other hand, results shown here are also germane to the debate on whether long-term data collection should continue to be a priority (if it actually ever was) in ecological research (Dupont et al. 2025; Blumstein 2025). Long-term pollinator *data* acquired in the field under the ecology-of-place paradigm (e.g., Janzen & Hallwachs 2021, present study) will not become redundant or irrelevant insofar as ecologists’ ingenuity succeeds in transforming them into *novel knowledge* helping to formulate and answer better ecological questions, understand the intricate action of the multiple drivers of population change, making predictions about the future, and last but not least fulfilling the ethical imperative of assembling an interpretable record of a vanishing world for the benefit of future generations ("Badía effect"; Pausas 2024).

## ACKNOWLEDGMENTS

I am deeply indebted to the many insect taxonomists that, by generously contributing identifications of specimens and photographs, made this study possible: Oscar Aguado (Apoidea, Symphyta), Jorge Almeida (Diptera), Miguel A. Alonso Zarazaga (Coleoptera), Piluca Álvarez (Diptera), Stig Andersen (Tachinidae), Rui Andrade (Diptera), Enrique Asensio (Apoidea), Marcos Báez (Bombyliidae), Leopoldo Castro (*Bombus*, Vespoidea), Antonio Cobos (Buprestidae), José García Carrillo (Anthicidae), Mario García-París (*Mylabris*), Severiano F. Gayubo (Crabronidae, Sphecidae), Max Kasparek (Anthidiini), María A. Marcos (Syrphidae), Andreas Müller (Megachilidae), Alejandro Núñez (Apoidea), Rafael Obregón (Coleoptera, Lepidoptera), Concepción Ornosa (Apidae), Francisco J. Ortiz-Sánchez (Apoidea), José Carlos Otero (Nitidulidae), Miguel Prieto (Dermestidae), Florent Prunier (Orthoptera), Antonio Ricarte (Syrphidae), Knut Rognes (Calliphoridae), Luis Rozas (Coleoptera, Neuroptera), Arabia Sánchez Terrón (Bombyliidae), Klaus Schönitzer (*Andrena*), Jan Smit (*Nomada*), Alberto Tinaut (Formicidae), Hans-Peter Tschorsnig (Tachinidae), Antonio Verdugo (*Anthaxia*), Amador Viñolas (*Aplocnemus*), Thomas J. Wood (*Andrena*) and José L. Yela (Lepidoptera). Alfredo Benavente and Gabriel Blanca assisted with some challenging plant identifications. Consejería de Medio Ambiente, Junta de Andalucía, granted permission to work in the Sierra de Cazorla and provided invaluable facilities there. Constructive comments and criticisms from Mark Genung and one anonymous reviewer were very helpful to refine and clarify this paper. Special thanks go to Mónica Medrano for insightful ideas, encouragement and inspiring discussions over so many years. The research reported in this paper received no specific grant from any funding agency.

## Data availability statement

Data files have been submitted to figshare (Herrera 2026) and are accessible to editors and reviewers via the following private link: https://figshare.com/s/b2c3fb7171d9ccec0eda

## Disclosure statement

The author declares no conflicts of interest.

## Generative AI disclosure statement

AI assistance was not used at any point in the preparation of this work.

## Appendices

Additional supporting information may be found in the online version of this article: Appendix 1: Additional figures and summaries of analytical results mentioned in the text.

## Notes

### Competing Interest Statement

The authors have declared no competing interest.

### Summary of Updates

Introduction was thoroughly re-written. New analyses and results were added. Two figures were dropped. New tables and figures added to the Appendix

## References

Adamo M, Chialva M, Calevo J, Bertoni F, Dixon K, Mammola S (2021) Plant scientists’ research attention is skewed towards colourful, conspicuous and broadly distributed flowers. Nature Plants 7:574–578.

Albrecht M, Schmid B, Hautier Y, Muller CB (2012) Diverse pollinator communities enhance plant reproductive success. Proceedings of the Royal Society B 279:4845–4852.

Alonso D, McKane AJ (2004) Sampling Hubbell’s neutral theory of biodiversity. Ecology Letters 7:901–910.

Alroy J (2015) The shape of terrestrial abundance distributions. Science Advances 1:e1500082.

Alroy J (2025) Does evenness even exist ? Ecology Letters 28:e70181.

Artamendi M, Martin PA, Bartomeus I, Magrach A (2025) Loss of pollinator diversity consistently reduces reproductive success for wild and cultivated plants. Nature Ecology & Evolution 9:296–313.

Austen GE, Bindemann M, Griffiths RA, Roberts DL (2016) Species identification by experts and non-experts: comparing images from field guides. Scientific Reports 6:33634.

Baldridge E, Harris DJ, Xiao X, White EP (2016) An extensive comparison of species-abundance distribution models. PeerJ 4:e2823.

Bates D, Mächler M, Bolker B, Walker S (2015) Fitting linear mixed-effects models using lme4. Journal of Statistical Software 67:1–48.

Beck JJ, Larget B, Waller DM (2018) Phantom species: adjusting estimates of colonization and extinction for pseudo-turnover. Oikos 127:1605–1618.

Blanco-Pastor JL, Ornosa C, Romero D, Liberal IM, Gómez JM, Vargas P (2015) Bees explain floral variation in a recent radiation of *Linaria*. Journal of Evolutionary Biology 28:851–863.

Blomberg SP, Garland T (2002) Tempo and mode in evolution: phylogenetic inertia, adaptation and comparative methods. Journal of Evolutionary Biology 15:899–910.

Blumstein DT (2025) The end of long-term ecological data? PLoS Biology 23:e3003102.

Blüthgen N (2010) Why network analysis is often disconnected from community ecology: A critique and an ecologist’s guide. Basic and Applied Ecology 11:185–195.

Blüthgen N, Klein AM (2011) Functional complementarity and specialisation: The role of biodiversity in plant–pollinator interactions. Basic and Applied Ecology 12:282–291.

Blüthgen N, Fründ J, Vázquez DP, Menzel F (2008) What do interaction network metrics tell us about specialization and biological traits? Ecology 89:3387–3399.

Bolker B, R Development Core Team (2023) bbmle: Tools for general maximum likelihood estimation. doi: 10.32614/CRAN.package.bbmle, R package version 1.0.25.1.

Burnham KP, Anderson DR (2002) Model selection and multimodel inference. Second edition. Springer, Berlin, Germany.

Chacoff NP, Resasco J, Vázquez DP (2018) Interaction frequency, network position, and the temporal persistence of interactions in a plant-pollinator network. Ecology 99:21–28.

Chao A, Jost L (2012) Coverage-based rarefaction and extrapolation: standardizing samples by completeness rather than size. Ecology 93:2533–2547.

Chao A, Jost L (2015) Estimating diversity and entropy profiles via discovery rates of new species. Methods in Ecology and Evolution 6:873–882.

Chao A, Gotelli NJ, Hsieh TC, Sander EL, Ma KH, Colwell RK, Ellison AM (2014) Rarefaction and extrapolation with Hill numbers: a framework for sampling and estimation in species diversity studies. Ecological Monographs 84:45–67.

Chao A, Colwell RK, Chiu CH, Townsend D (2017) Seen once or more than once: applying Good-Turing theory to estimate species richness using only unique observations and a species list. Methods in Ecology and Evolution 8:1221–1232.

Corbet AS (1941) The distribution of butterflies in the Malay Peninsula. Proceedings of the Royal Entomological Society of London A 16:101–116.

Davila YC, Elle E, Vamosi JC, Hermanutz L, Kerr JT, Lortie CJ, Westwood AR, Woodcock TS, Worley AC (2012) Ecosystem services of pollinator diversity: a review of the relationship with pollen limitation of plant reproduction. Botany 90:535–543.

Dormann CF, Gruber B, Fründ J (2008) Introducing the bipartite package: analysing ecological networks. R News 8:8–11.

Doyle T, Hawkes WLS, Massy R, Powney GD, Menz MHM, Wotton KR (2020) Pollination by hoverflies in the Anthropocene. Proceedings of the Royal Society B 287:20200508.

Dupont L, Jacob S, Philippe H (2025) Scientist engagement and the knowledge–action gap. Nature Ecology & Evolution 9:23–33.

Fisher RA, Corbet AS, Williams CB (1943) The relation between the number of species and the number of individuals in a random sample of an animal population. Journal of Animal Ecology 12:42–58.

Fründ J, Linsenmair KE, Blüthgen N (2010) Pollinator diversity and specialization in relation to flower diversity. Oikos 119:1581–1590.

Fründ J, Dormann CF, Holzschuh A, Tscharntke T (2013) Bee diversity effects on pollination depend on functional complementarity and niche shifts. Ecology 94:2042–2054.

Gallagher MK, Campbell DR (2017) Shifts in water availability mediate plant-pollinator interactions. New Phytologist 215:792–802.

Galloni M, Podda L, Vivarelli D, Quaranta M, Cristofolini G (2008) Visitor diversity and pollinator specialization in Mediterranean legumes. Flora 203:94–102.

Garcia JE, Hannah L, Shrestha M, Burd M, Dyer AG (2022) Fly pollination drives convergence of flower coloration. New Phytologist 233:52–61.

Genung MA, Fox J, Willms NM, Kremen C, Ascher J, Gibbs J, Winfree R (2017) The relative importance of pollinator abundance and species richness for the temporal variance of pollination services. Ecology 98:1807–1816.

Genung MA, Reilly J, Williams NM, Buderi A, Gardner J, Winfree R (2023) Rare and declining bee species are key to consistent pollination of wildflowers and crops across large spatial scales. Ecology 104:e3899.

Gómez JM, Bosch J, Perfectti F, Fernández J, Abdelaziz M (2007) Pollinator diversity affects plant reproduction and recruitment: the tradeoffs of generalization. Oecologia 153:597–605.

Gómez JM, Perfectti F, Abdelaziz M, Lorite J, Muñoz-Pajares AJ, Valverde J (2015) Evolution of pollination niches in a generalist plant clade. New Phytologist 205:440–453.

Gómez Mercado F (2011) Vegetación y flora de la Sierra de Cazorla. Guineana 17:1–481.

Gong YB, Huang SQ (2009) Floral symmetry: pollinator-mediated stabilizing selection on flower size in bilateral species. Proceedings of the Royal Society B 276:4013–4020.

Gotelli NJ, Chao A (2013) Measuring and estimating species richness, species diversity, and biotic similarity from sampling data. Encyclopedia of Biodiversity 5:195–211.

Hall M (2018) Blue and yellow vane traps differ in their sampling effectiveness for wild bees in both open and wooded habitats. Agricultural and Forest Entomology 20:487–495.

Hartigan JA, Hartigan PM (1985) The dip test of unimodality. Annals of Statistics 13:70–84.

Herrera CM (1987a) Components of pollinator ’quality’: comparative analysis of a diverse insect assemblage. Oikos 50:79–90.

Herrera CM (1987b) Componentes del flujo génico en *Lavandula latifolia* Medicus: polinización y dispersión de semillas. Anales del Jardín Botánico de Madrid 44:49–61.

Herrera CM (1989) Pollinator abundance, morphology, and flower visitation rate: analysis of the ’quantity’ component in a plant-pollinator system. Oecologia 80:241–248.

Herrera CM (1990) Daily patterns of pollinator activity, differential pollinating effectiveness, and floral resource availability, in a summer-flowering Mediterranean shrub. Oikos 58:277–288.

Herrera CM (1995) Floral biology, microclimate, and pollination by ectothermic bees in an early-blooming herb. Ecology 76:218–228.

Herrera CM (2000) Flower-to-seedling consequences of different pollination regimes in an insect-pollinated shrub. Ecology 81:15–29.

Herrera CM (2005) Plant generalization on pollinators: species property or local phenomenon? American Journal of Botany 92:13–20.

Herrera CM (2019) Complex long-term dynamics of pollinator abundance in undisturbed Mediterranean montane habitats over two decades. Ecological Monographs 89:e01338.

Herrera CM (2020) Flower traits, habitat, and phylogeny as predictors of pollinator service: a plant community perspective. Ecological Monographs 90:e01402.

Herrera CM (2021) Unclusterable, underdispersed arrangement of insect-pollinated plants in pollinator niche space. Ecology 102:e03327.

Herrera CM (2024) Thermal biology diversity of bee pollinators: taxonomic, phylogenetic and plant community-level correlates. Ecological Monographs 94:e1625.

Herrera CM (2026) Data: Species abundance and diversity of a Mediterranean insect pollinator assemblage. Data. Figshare. https://figshare.com/s/b2c3fb7171d9ccec0eda

Herrera CM, Núñez A, Valverde J, Alonso C (2023) Body mass decline in a Mediterranean community of solitary bees supports the size shrinking effect of climatic warming. Ecology 104:e4128.

Herrera CM, Alonso C, Valverde J, Núñez A, Wood TJ (2025) The genus *Andrena* Fabricius, 1775 (Hymenoptera, Andrenidae) in a Mediterranean biodiversity hotspot: community-wide relationships with plants and description of three new species. Journal of Hymenoptera Research 98:1039–1066.

Hill MO (1973) Diversity and evenness: a unifying notation and its consequences. Ecology 54:427–432.

Hsieh TC, Ma KH, Chao A (2016) iNEXT: an R package for rarefaction and extrapolation of species diversity (Hill numbers). Methods in Ecology and Evolution 7:1451–1456.

Hu L, Dong Y, Sun S (2019) Relative species abundance successfully predicts nestedness and interaction frequency of monthly pollination networks in an alpine meadow. PLoS ONE 14:e0224316.

Hubbell SP (2001) The unified neutral theory of biodiversity and biogeography. Princeton University Press, Princeton, NJ, USA.

Janzen DH, Hallwachs W (2021) To us insectometers, it is clear that insect decline in our Costa Rican tropics is real, so let’s be kind to the survivors. Proceedings of the National Academy of Sciences 118:e2002546117.

Jin Y, Qian H (2022) V.PhyloMaker2: An updated and enlarged R package that can generate very large phylogenies for vascular plants. Plant Diversity 44:335–339.

Jost L (2006) Entropy and diversity. Oikos 113:363–375.

Jost L (2010) The relation between evenness and diversity. Diversity 2:207–232.

Kalin Arroyo MT, Primack R, Armesto J (1982) Community studies in pollination ecology in the high temperate Andes of central Chile. I. Pollination mechanisms and altitudinal variation. American Journal of Botany 69:82–97.

Kato M, Kakutani T, Inoue T, Itino T (1990) Insect-flower relationship in the primary beech forest of Ashu, Kyoto: An overview of the flowering phenology and the seasonal pattern of insect visits. Contributions Biological Laboratory Kyoto University 27:309–375

Keck F, Rimet F, Bouchez A, Franc A (2016) phylosignal: an R package to measure, test, and explore the phylogenetic signal. Ecology and Evolution 6:2774–2780.

Kempton RA, Taylor LR (1974) Log-series and log-normal parameters as diversity discriminants for the Lepidoptera. Journal of Animal Ecology 43:381–399.

Kevan PG, Greco CF, Belaoussoff S (1997) Log-normality of biodiversity and abundance in diagnosis and measuring of ecosystemic health: pesticide stress on pollinators on blueberry heaths. Journal of Applied Ecology 34:1122–1136.

King C, Ballantyne G, Willmer PG (2013) Why flower visitation is a poor proxy for pollination: measuring single-visit pollen deposition, with implications for pollination networks and conservation. Methods in Ecology and Evolution 4:811–818.

Knuth P (1908) Handbook of flower pollination. Vol. 2. Translated by J. R. Ainsworth Davis. Clarendon, Oxford, UK.

Lanuza JB, Knight TM, Montes-Perez N, Glenny W, Acuña P, Albrecht M, Artamendi M, Badenhausser I, Bennett JM, Biella P, et al (2025) EuPPollNet: A European database of plant-pollinator networks. Global Ecology and Biogeography 34:e70000.

Larsen S, Chase JM, Durance I, Ormerod SJ (2018) Lifting the veil: richness measures fail to detect systematic biodiversity change over three decades. Ecology 99:1316–1326.

Liu J, Slik F, Zheng S, Lindenmayer DB (2022) Undescribed species have higher extinction risk than known species. Conservation Letters 15:e12876.

Lüdecke D (2018) ggeffects: Tidy data frames of marginal effects from regression models. Journal of Open Source Software 3:772.

Lüttge U (2013) Green nectaries: the role of photosynthesis in secretion. Botanical Journal of the Linnean Society 173:1–11.

Loy X, Brosi BJ (2022) The effects of pollinator diversity on pollination function. Ecology 103:e3631.

Magurran AE (2004) Measuring biological diversity. Blackwell, Oxford, UK.

Margalef R (1957) La teoría de la información en ecología. Memorias de la Real Academia de Ciencias y Artes de Barcelona 32:373–449.

Margalef R (1968) Perspectives in ecological theory. University of Chicago Press, Chicago, IL, USA.

Marini L, Gazzea E, Albrecht M, Báldi A, Batáry P, Bartomeus I, Bommarco R, Bruun HH, Cappellari A, Cole LJ, et al (2025) Using total abundance as a proxy for wild bee species richness: A practical tool for non-[experts. Journal of Applied Ecology 62:3065–3077.

Martins KT, Gonzalez A, Lechowicz MJ (2015) Pollination services are mediated by bee functional diversity and landscape context. Agriculture, Ecosystems & Environment 200:12–20.

Matthews TJ, Whittaker RJ (2015) On the species abundance distribution in applied ecology and biodiversity management. Journal of Applied Ecology 52:443–454.

May RM (1975) Patterns of species abundance and diversity. Pages 81–120 in M. L. Cody and J. M. Diamond, editors. Ecology and evolution of communities. Belknap Press, Cambridge, MA, USA.

Maynard DS, Servan CA, Allesina S (2018) Network spandrels reflect ecological assembly. Ecology Letters 21:324–334.

McGill BJ, Etienne RS, Gray JS, Alonso D, Anderson MJ, Benecha HK, Dornelas M, Enquist BJ, Green JL, He F (2007) Species abundance distributions: moving beyond single prediction theories to integration within an ecological framework. Ecology Letters 10:995–1015.

Médail F, Diadema K (2009) Glacial refugia influence plant diversity patterns in the Mediterranean Basin. Journal of Biogeography 36:1333–1345.

Moldoveanu OC, Maggioni M, Vergari D, Dani FR (2025) Revealing the diversity of wild bees (Hymenoptera: Apoidea: Anthophila) in flower strips in Mediterranean floodplains: which monitoring method fits best? Biodiversity and Conservation 34:2809–2828.

Montgomery BR, Rathcke BJ (2012) Effects of floral restrictiveness and stigma size on heterospecific pollen receipt in a prairie community. Oecologia 168:449–458.

Müller H (1883) The fertilisation of flowers. Translated by D. W. Thompson. MacMillan, London, UK.

Münkemüller T, Lavergne S, Bzeznik B, Dray S, Jombart T, Schiffers K, Thuiller W (2012) How to measure and test phylogenetic signal. Methods in Ecology and Evolution 3:743–756.

Norfolk O, Eichhorn MP, Gilbert FS (2015) Contrasting patterns of turnover between plants, pollinators and their interactions. Diversity and Distributions 21:405–415.

Ollerton J (2017) Pollinator diversity: distribution, ecological function, and conservation. Annual Review of Ecology, Evolution, and Systematics 48:353–376.

Ollerton J, Winfree R, Tarrant S (2011) How many flowering plants are pollinated by animals? Oikos 120:321–326.

Ollerton J, Rech AR, Waser NM, Price MV (2015) Using the literature to test pollination syndromes – some methodological cautions. Journal of Pollination Ecology 16:119–125.

Ortiz-Sánchez FJ, Valverde J, Núñez Carbajal A, Alonso C, Herrera CM (2023) The bees (Hymenoptera, Apoidea) of Sierra de Cazorla (Spain). Monografías de la Sociedad Entomológica Aragonesa 17:1–91.

Packer L, Monckton SK, Onuferko TM, Ferrari RR (2018) Validating taxonomic identifications in entomological research. Insect Conservation and Diversity 11:1–12.

Pärtel M, Szava-Kovats R, Zobel M (2011) Dark diversity: shedding light on absent species. Trends in Ecology and Evolution 26:124–128.

Pausas JG (2024) Science in a changing world. Frontiers in Ecology and the Environment 22:e2797.

Peet RK (1974) The measurement of species diversity. Annual Review of Ecology and Systematics 5:285–307.

Pérez-Gómez Á, Godoy O, Ojeda F (2024) Beware of trees: pine afforestation of a naturally treeless habitat reduces flower and pollinator diversity. Global Ecology and Conservation 50:e02808.

Pérez-Méndez N, Andersson GKS, Requier F, Hipólito J, Aizen MA, Morales CL, García N, Gennari GP, Garibaldi LA (2020) The economic cost of losing native pollinator species for orchard production. Journal of Applied Ecology 57:599–608.

Petanidou T (1991) Pollinating fauna of a phryganic ecosystem: species list. Verslagen en technische gegevens, Instituut voor Taxonomische Zoölogie (Zoölogisch Museum) Universiteit van Amsterdam 59:1–11.

Petanidou T, Ellis WN (1993) Pollinating fauna of a phryganic ecosystem: composition and diversity. Biodiversity Letters 1:9–22.

Prado PI, Miranda MD, Chalom A (2025) sads: Maximum likelihood models for species abundance distributions. https://CRAN.R-project.org/package=sads

Price MV, Billick I, editors (2010) The ecology of place. University of Chicago Press, Chicago, IL, USA.

Pugnaire FI, Arista M, Carrión JS, Devesa JA, Herrera CM, Nieto Feliner G, Rey PJ, Alonso C (2024) Origen y retos actuales de la diversidad vegetal en las sierras Béticas, un área de diversidad de importancia global. Ecosistemas 33:2676.

R Core Team (2025) R: A language and environment for statistical computing. R Foundation for Statistical Computing, Vienna, Austria.

Ratnieks FLW, Schrell F, Sheppard RC, Brown E, Bristow OE, Garbuzov M (2016) Data reliability in citizen science: learning curve and the effects of training method, volunteer background and experience on identification accuracy of insects visiting ivy flowers. Methods in Ecology and Evolution 7:1226–1235.

Renaud E, Baudry E, Bessa-Gomes C (2020) Influence of taxonomic resolution on mutualistic network properties. Ecology and Evolution 10:3248–3259.

Rezende SM, Pennisi SV, Gariepy T, Querejeta M, Ulyshen M, Schmidt JM (2025) Wild bees show local spatial and temporal dynamics in southeastern US blueberry farmscapes. Environmental Entomology 54:67–76.

Ricotta C, Feoli E (2024) Hill numbers everywhere. Does it make ecological sense? Ecological Indicators 161:111971.

Robertson C (1928) Flowers and insects. Lists of visitors of four hundred and fifty-three flowers. Science Press, Lancaster, Pennsylvania, USA.

Roswell M, Dushoff J, Winfree R (2021) A conceptual guide to measuring species diversity. Oikos 130:321–338.

Roswell M, Harrison T, Genung MA (2023) Biodiversity–ecosystem function relationships change in sign and magnitude across the Hill diversity spectrum. Philosophical Transactions of the Royal Society B 378:20220186.

Serra-Marin PE, Solé-Ribalta A, Lana A, Borge-Holthoefer J, Hervías-Parejo S, Traveset A (2025) Comparative assessment of automated and manual monitoring in comprehensive plant–pollinator communities. Methods in Ecology and Evolution 16:2960–2978.

Simpson DT, Weinman LR, Genung MA, Roswell M, MacLeod M, Winfree R (2022) Many bee species, including rare species, are important for function of entire plant-pollinator networks. Proceedings of the Royal Society B 289:20212689.

Su Q (2018) A general pattern of the species abundance distribution. PeerJ 6:e5928.

Taylor SJS, Evans BS, White EP, Hurlbert AH (2018) The prevalence and impact of transient species in ecological communities. Ecology 99:1825–1835.

Tepedino VJ, Stanton NL (1981) Diversity and competition in bee-plant communities on short-grass prairie. Oikos 36:35–44.

Thompson A, Frenzel M, Schweiger O, Musche M, Groth T, Roberts SPM, Kuhlmann M, Knight TM (2021) Pollinator sampling methods influence community patterns assessments by capturing species with different traits and at different abundances. Ecological Indicators 132:108284.

Thomson JD (2021) How worthwhile are pollination networks? Journal of Pollination Ecology 28. Doi: 10.26786/1920-7603(2021)652

Ulrich W, Ollik M, Ugland KI (2010) A meta-analysis of species-abundance distributions. Oikos 119:1149–1155.

Valverde J, Perfectti F, Gómez JM (2019) Pollination effectiveness in a generalist plant: adding the genetic component. New Phytologist 223:354–365.

Vázquez DP, Blüthgen N, Cagnolo L, Chacoff NP (2009) Uniting pattern and process in plant-animal mutualistic networks: a review. Annals of Botany 103:1445–1457.

Venjakob C, Klein AM, Ebeling A, Tscharntke T, Scherber C (2016) Plant diversity increases spatio-temporal niche complementarity in plant-pollinator interactions. Ecology and Evolution 6:2249–2261.

Waser NM, Price MV (2016) Drought, pollen and nectar availability, and pollination success. Ecology 97:1400–1409.

Watazu T, Hiraiwa MK, Inoue M, Mishima H, Ushimaru A, Hosaka T (2025) Effectiveness of interval photography cameras for a survey of pollinator communities: Comparison with direct observation. Applications in Plant Sciences 13:e70023.

Whittaker RH (1965) Dominance and diversity in land plant communities. Science 147:250–260.

Williams CB (1964) Patterns in the balance of nature and related problems in quantitative ecology. Academic Press, London, UK.

Winfree R, Williams NM, Dushoff J, Kremen C (2014) Species abundance, not diet breadth, drives the persistence of the most linked pollinators as plant-pollinator networks disassemble. American Naturalist 183:600–611.

Winfree R, Fox JW, Williams NM, Reilly JR, Cariveau DP (2015) Abundance of common species, not species richness, drives delivery of a real-world ecosystem service. Ecology Letters 18:626–635.

Wood SN (2017) Generalized additive models: An introduction with R. 2nd edition. CRC Press, Boca Raton, FL, USA.

Wood S, Scheipl F (2025) gamm4: Generalized Additive Mixed Models using mgcv and lme4. doi: 10.32614/CRAN.package.gamm4

