## Appendix 1 for "Commonness and rarity of pollinators: species abundance and diversity in a hyperdiverse Mediterranean assemblage"

Appendix 1. Additional figures and summaries of analytical results mentioned in the text.

Figure 1. Bivariate plots depicting the relationship between the rank abundances of individual pollinator species based on total number of individuals recorded and total number of flowers visited, at the regional (left) and local (right) spatial scales. Each green circle represents one pollinator species (*N* = 845 and 583 for regional and local scales, respectively), red lines are least squares-fitted linear regressions (*R*^2^ = 0.91 and 0.87 for regional and local analyses, respectively), and dashed lines are *y* = *x* isolines.

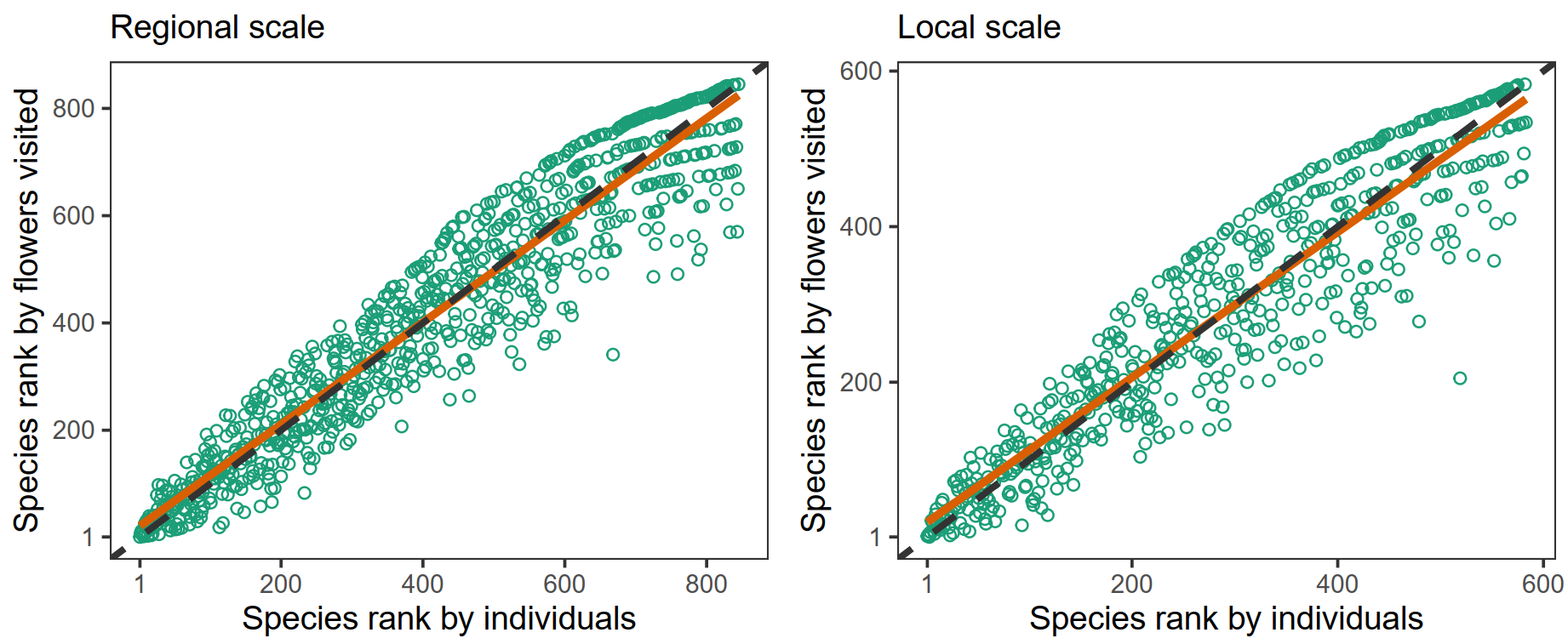

Figure 2. Relationships between diversity measures and estimated sample coverage (SC) in the set of 447 pollinator sampling occasions ( = plant species x site x year combinations) on the 275 plant species included in the study. Each symbol is a sampling occasion and lines are ordinary least squares-fitted regressions.

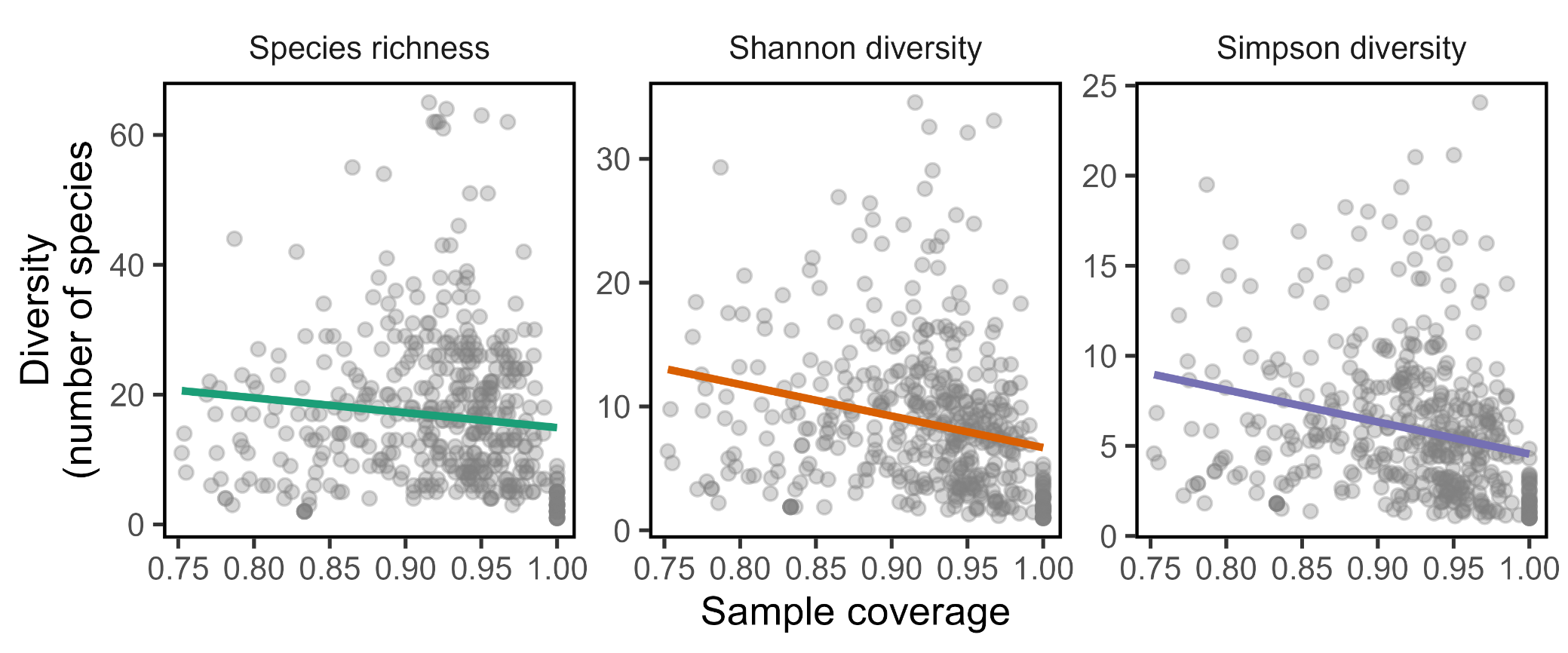

| Figure 3. Phylogenetic relationships of the *N* = 275 plant species included in the diversity analyses, and variation across plant species in Shannon and Simpson pollinator diversity measures. Tree tips are color-coded to denote whether the pollinator diversity estimate of the plant species involved falls above or below the median of the overall distribution for that particular diversity measurement (red and grey dots, respectively). Selected clades exemplifying predominantly high (Asteraceae, Convolvulaceae, Rosaceae) or low (Brassicaceae, Fabaceae, Plantaginaceae) pollinator diversity are highlighted (red and grey sectors, respectively). | |
| --- | --- |
| **Shannon diversity**  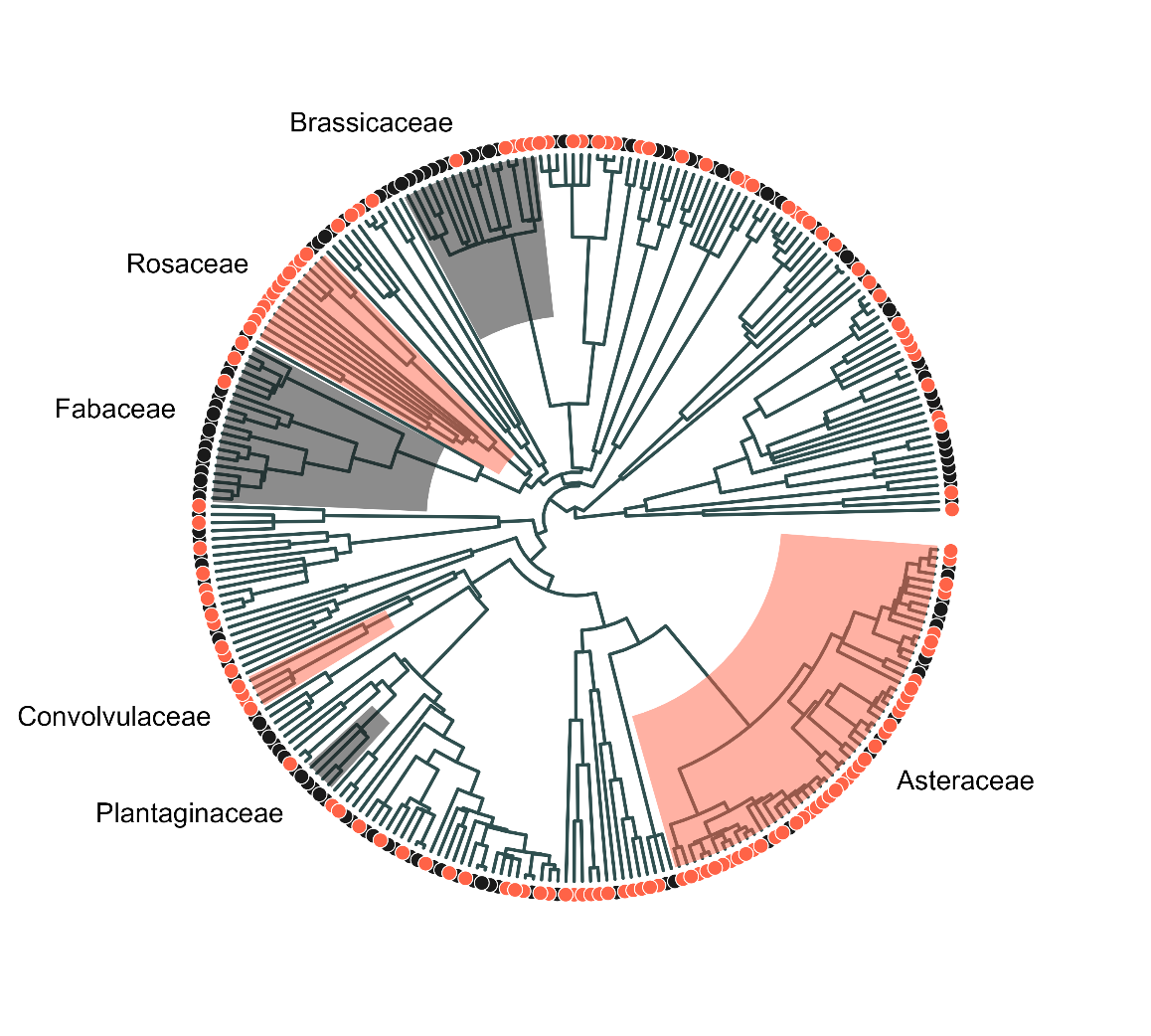 | **Simpson diversity**  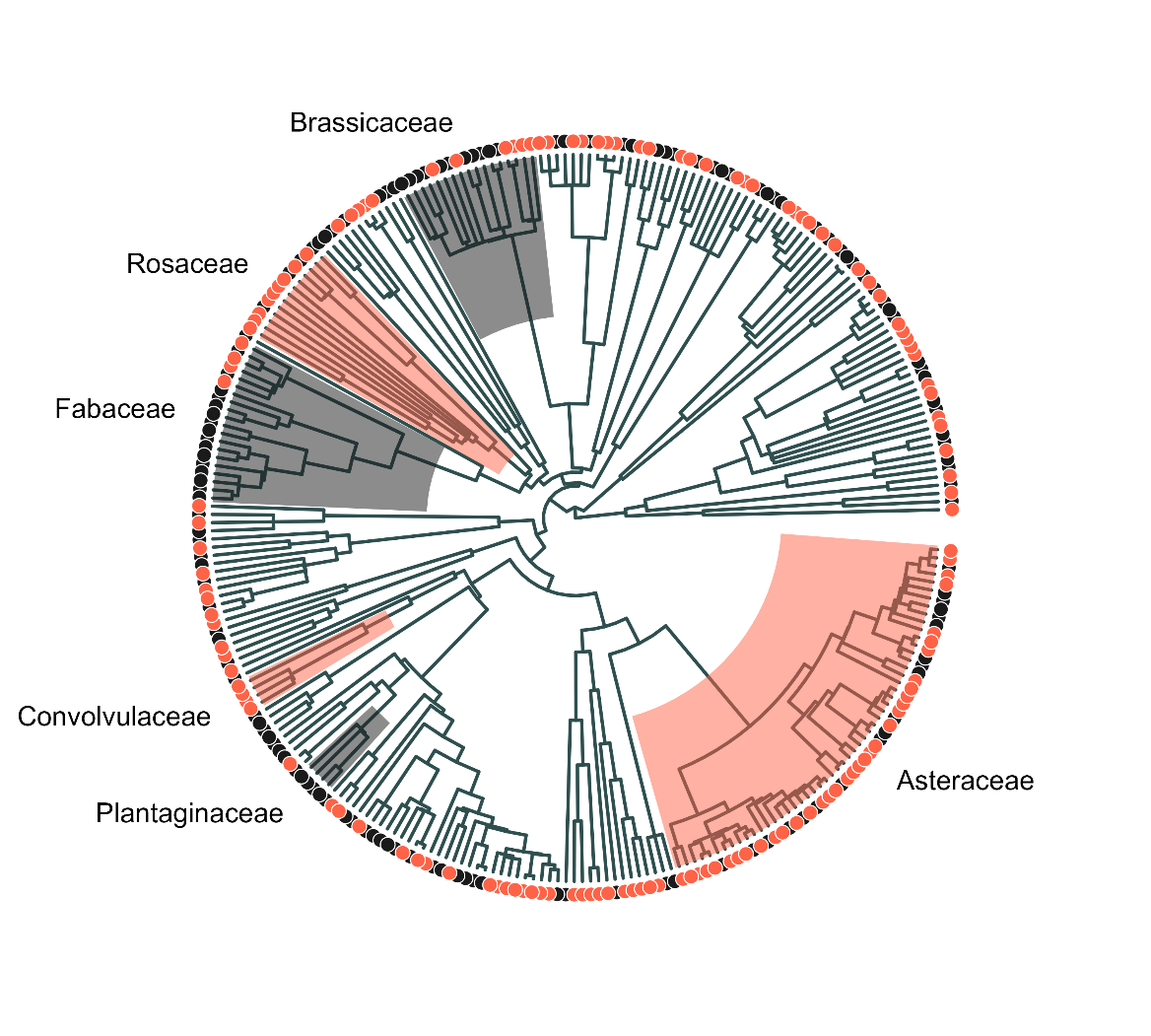 |

| Figure 4. Interaction plots showing model-adjusted, estimated marginal means (dots, vertical segments are 95% confidence intervals) of Shannon (left) and Simpson (right) pollinator diversity for plant species differing in perianth type (open vs. restrictive) and pollinator visitation unit (single flower vs. flower packet). See Table 3 in this Appendix for model details and analytical results. | |
| --- | --- |
| 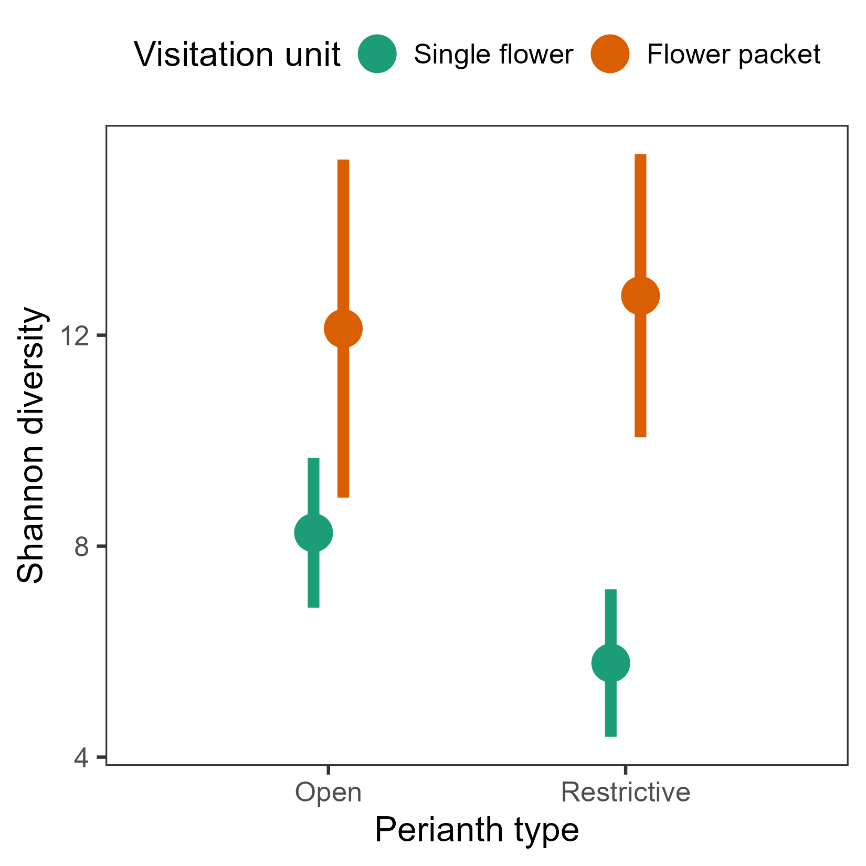 | 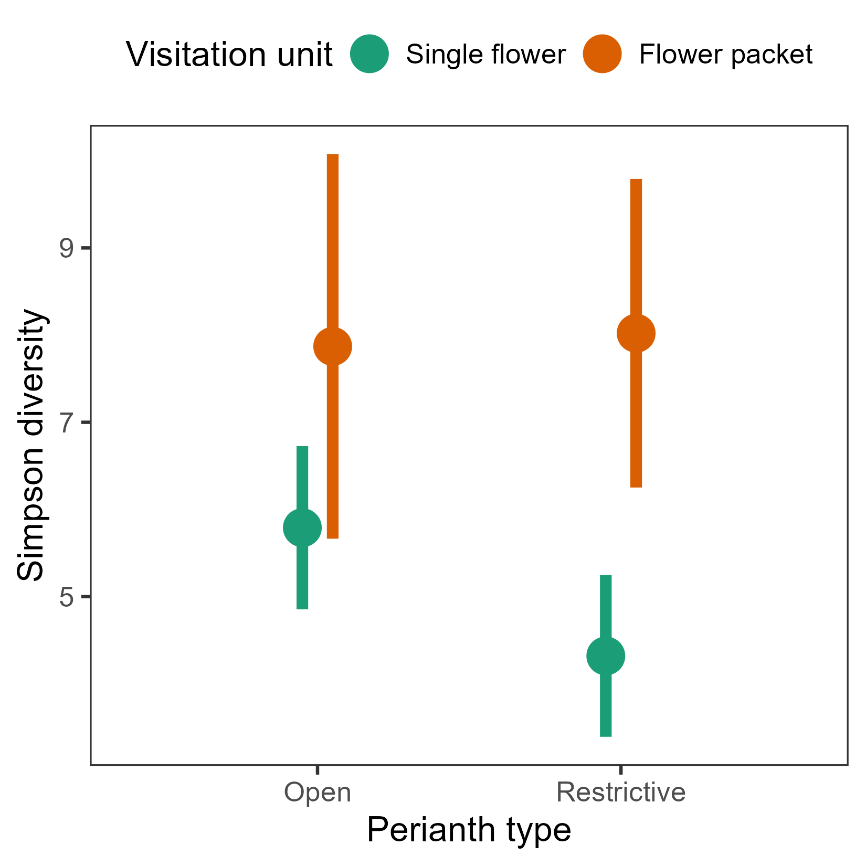 |

Table 1 Information criteria to compare the performance of five statistical distributions to predict empirical species abundance distributions of the four major insect orders. For each order models are listed in decreasing order of performance. AIC, Akaike's Information Criterion; Δ, AIC difference, or the increment of AIC relative to the minimum value; df = degrees of freedom, equal to the number of free parameters in each statistical model; *w* = Akaike's weight, or the relative likelihood of the model given the data, normalized to sum to 1 and interpretable as probabilities.

| Insect order | Species | Individuals | Statistical model | AIC | Δ | df | *w* |
| --- | --- | --- | --- | --- | --- | --- | --- |
| Coleoptera: | 128 | 8,249 |  |  |  |  |  |
|  |  |  | Log-series | 1075.2 | 0.0 | 1 | 0.45 |
|  |  |  | Neutral metacommunity | 1075.4 | 0.1 | 1 | 0.42 |
|  |  |  | Poisson lognormal | 1077.8 | 2.6 | 2 | 0.12 |
|  |  |  | Power law | 1094.0 | 18.7 | 1 | <0.001 |
|  |  |  | Negative binomial | 1337.4 | 262.1 | 2 | <0.001 |
| Diptera: | 235 | 9,622 |  |  |  |  |  |
|  |  |  | Log-series | 1911.4 | 0.0 | 1 | 0.58 |
|  |  |  | Neutral metacommunity | 1912.2 | 0.8 | 1 | 0.40 |
|  |  |  | Poisson lognormal | 1918.4 | 7.0 | 2 | 0.02 |
|  |  |  | Power law | 1974.7 | 63.3 | 1 | <0.001 |
|  |  |  | Negative binomial | 2421.0 | 509.6 | 2 | <0.001 |
| Hymenoptera: | 349 | 22,111 |  |  |  |  |  |
|  |  |  | Log-series | 3132.9 | 0.0 | 1 | 0.55 |
|  |  |  | Neutral metacommunity | 3133.3 | 0.4 | 1 | 0.44 |
|  |  |  | Poisson lognormal | 3144.2 | 11.4 | 2 | 0.002 |
|  |  |  | Power law | 3257.1 | 124.2 | 1 | <0.001 |
|  |  |  | Negative binomial | 3890.5 | 757.6 | 2 | <0.001 |
| Lepidoptera | 108 | 6,099 |  |  |  |  |  |
|  |  |  | Neutral metacommunity | 966.2 | 0.0 | 1 | 0.52 |
|  |  |  | Log-series | 966.4 | 0.2 | 1 | 0.48 |
|  |  |  | Poisson lognormal | 977.0 | 10.9 | 2 | 0.002 |
|  |  |  | Power law | 1008.9 | 42.8 | 1 | <0.001 |
|  |  |  | Negative binomial | 1170.0 | 203.9 | 2 | <0.001 |

Table 2. Summary statistics of distributions of plant species means (*N* = 275 plant species) shown in Figure 4 for the three pollinator diversity measures considered. Diversities are expressed in number of species.

| Diversity measure | Range | Interquartile range | Median | Mean (SE) |
| --- | --- | --- | --- | --- |
| Species richness | 1-62 | 7-21 | 13 | 15.5 (0.65) |
| Shannon diversity | 1-30.8 | 4.0-10.8 | 7.4 | 8.1 (0.32) |
| Simpson diversity | 1-20.3 | 2.9-7.6 | 4.8 | 5.6 (0.22) |

Table 3. Summary of results of the three linear mixed-effects models fitted to test for effects of floral features (perianth type, visitation unit, and their interaction) on plant species means for each pollinator diversity measurement.

|  | Effect | | | | | | | |
| --- | --- | --- | --- | --- | --- | --- | --- | --- |
|  | Perianth type (PT) | |  | Visitation unit (VU) | |  | PT x VU | |
| Response variable | Chi-square | *P*-value |  | Chi-square | *P*-value |  | Chi-square | *P*-value |
| Species richness | 9.4 | 0.0021 |  | 36.7 | 1.4E-09 |  | 2.2 | 0.14 |
| Shannon diversity | 5.4 | 0.019 |  | 23.8 | 1.1E-06 |  | 1.9 | 0.16 |
| Simpson diversity | 4.4 | 0.035 |  | 15.2 | 9.7E-05 |  | 1.2 | 0.28 |

*Notes*: Plant family was included as a random effect in all models to account for family-level phylogenetic signal. Results of the random part of models are omitted from the table. R syntax for the mixed-effects models fitted to estimate the effects of floral features and their interaction on diversity measurements: lmer(diversity ~ perianth.type + visitation.unit + visitation.unit:perianth.type + (1|plant.family). There was no variance in floral traits within most plant genera, so plant genus was not included among random effects.

Table 4. Summary of the nonparametric regressions shown in Figure 6 to depict the relationships between two pollinator visitation measurements (visitation probability per time unit to flowering patches and to individual flowers, all pollinator species combined) and three pollinator diversity measures (Species richness, Shannon diversity and Simpson diversity) in the set of 275 plant species studied.

|  |  |  |  | Smooth term | | |
| --- | --- | --- | --- | --- | --- | --- |
| Response variable | Predictor | % deviance explained |  | edf | *F* | *P*-value |
| Patch visitation probability | ^0^D | 74.6 |  | 3.85 | 183.5 | <2E-16 |
|  | ^1^D | 49.1 |  | 3.06 | 65.7 | <2E-16 |
|  | ^2^D | 35.8 |  | 3.09 | 35.9 | <2E-16 |
| Flower visitation probability | ^0^D | 17.2 |  | 3.02 | 15.7 | <2E-16 |
|  | ^1^D | 19.0 |  | 2.88 | 13.5 | <2E-16 |
|  | ^2^D | 14.4 |  | 3.31 | 7.7 | 3.4E-05 |

*Notes:* Species richness, Shannon diversity and Simpson diversity of pollinators are coded as ^0^D, ^1^D and ^2^D, respectively. Regressions were fitted to plant species means for each predictor-response pair, and species means for sample coverage (SC) were also included as a fixed-effect covariate to statistically control for the (weak) relationship between SC and diversity (Figure 2 in this Appendix). edf = estimated degrees of freedom of the fitted smoothing spline. R syntax for nonparametric regressions: gam(visitation.probability ~ sample.coverage + s(Diversity, bs = "cr", k = 5)

Table 5. Summary of results from fitting separate linear mixed-effects models to plant species means for the three pollinator diversity measures (response variable), testing for the effect of habitat type (9-level, fixed-effect factor) while statistically accounting for differences among habitats in taxonomic composition of plant communities by including plant family, and genus nested within family, as random effects.

|  | Model *R*^2^ | |  | Fixed effect (Habitat type) | | |  | Estimated variance of random effects | |
| --- | --- | --- | --- | --- | --- | --- | --- | --- | --- |
| Response variable | Marginal (fixed effect only) | Conditional (fixed + random effects) |  | Chi-square | df | *P*-value |  | Plant family | Plant genus within family |
| Species richness | 0.116 | 0.641 |  | 63.7 | 8 | 8.6E-11 |  | 56.8 | 10.9 |
| Shannon diversity | 0.100 | 0.506 |  | 43.4 | 8 | 7.3E-07 |  | 11.1 | 2.0 |
| Simpson diversity | 0.083 | 0.399 |  | 30.4 | 8 | 0.00018 |  | 3.3 | 1.2 |

*Notes*: R syntax for mixed-effects models fitted: lmer(Diversity ~ habitat.type + (1|plant.family/genus).

Table 6. Summary of results of generalized additive mixed-effects models testing for seasonal (within year) and long-term (among years) trends in pollinator diversity.

|  |  |  |  | Smooth term | | |
| --- | --- | --- | --- | --- | --- | --- |
| Response | Model *R*^2^ | Predictor |  | edf | *F* | *P*-value |
| Species richness | 0.223 | Time of year |  | 3.62 | 12.7 | <2.0E-16 |
|  |  | Year |  | 6.11 | 5.9 | 6.0E-06 |
| Shannon diversity | 0.193 | Time of year |  | 3.40 | 11.7 | 2.2E-07 |
|  |  | Year |  | 4.98 | 3.0 | 0.012 |
| Simpson diversity | 0.142 | Time of year |  | 3.17 | 9.7 | 3.0E-06 |
|  |  | Year |  | 2.87 | 2.5 | 0.099 |

*Notes:* Models were fitted to the pollinator data from the *N* = 447 sampling occasions (= plant species x year x site combinations). Models included plant species and sampling site as random effects (results of the random part of the models are omitted from the table). edf = estimated degrees of freedom of the fitted smoothing spline. R syntax for mixed-effects models fitted: gamm4(Diversity ~ s(mean.julian.date, bs = "cr", k = -1) + s(year, bs = "cr", k = -1), random = ~ (1|plant.species) + (1|sitio).
